# Ensemble and Iterative Engineering for Maximized Bioconversion to the Blue Pigment, Indigoidine from Non-Canonical Sustainable Carbon Sources

**DOI:** 10.1101/2023.03.16.532821

**Authors:** Thomas Eng, Deepanwita Banerjee, Javier Menasalvas, Yan Chen, Jennifer Gin, Hemant Choudhary, Edward Baidoo, Jian Hua Chen, Axel Ekman, Ramu Kakumanu, Yuzhong Liu Diercks, Alex Codik, Carolyn Larabell, John Gladden, Blake Simmons, Jay D. Keasling, Christopher J. Petzold, Aindrila Mukhopadhyay

## Abstract

While many heterologous molecules can be produced via microbial bioconversion processes, maximizing their titers, rates, and yields from lignin-derived carbon streams remains challenging. Growth coupling can not only increase titers and yields but also shift the production period from stationary phase to growth phase. These methods for designing growth-coupling strains however require multi-gene edits for implementation which may be perceived as impractical. Here, we computationally evaluated 4,114 potential solutions for growth coupling *para-*coumarate to indigoidine production and prototype two cut sets in *Pseudomonas putida* KT2440. We used adaptive laboratory evolution (ALE) on the initial triple deletion strain to restore growth on *p-*CA. Using X-ray tomography on this post-ALE strain we revealed increased cell density and decreased cell volume. Proteomics identified upregulated peroxidases that mitigate reactive oxygen species formation. Nine iterative stepwise modifications further informed by model-guided and rational approaches realized a growth coupled strain that produced 7.3 g/L indigoidine at 77% MTY in *para*-coumarate minimal medium. These ensemble strategies provide a blueprint for producing target molecules at high product titers, rates, and yields.

**GRAPHICAL ABSTRACT:** 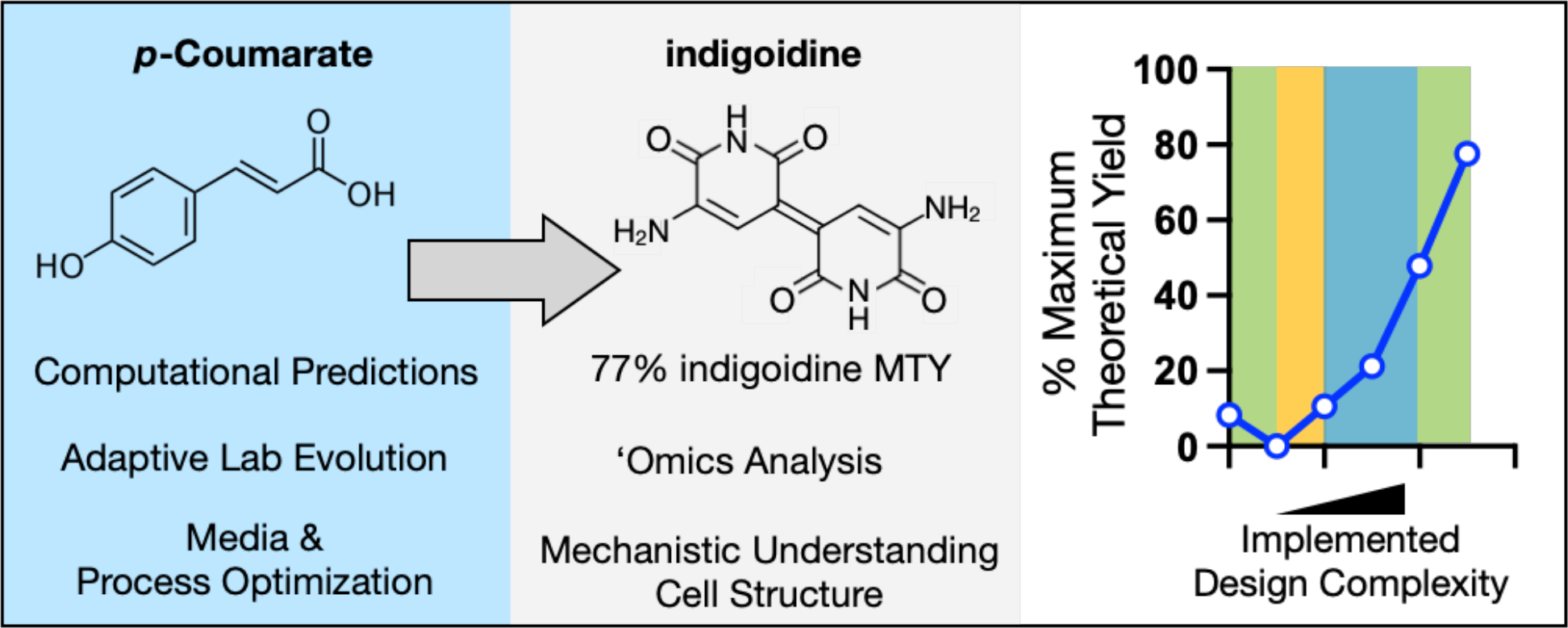

## INTRODUCTION

Microbial genomes encode an underexplored reservoir of natural products that have applications in health and biomanufacturing (Skinnider *et al*, 2020; Beller *et al*, 2015; Skoog *et al*, 2018; Keasling *et al*, 2021). It is well established that proof of concept quantities of many compounds can be expressed in genetically tractable hosts, but increasing their titers, rates, and yields to appreciable levels is a crucial step necessary for deployment at commercial scales (Blöbaum *et al*, 2023; Wehrs *et al*, 2019b). When calculating the economic viability of any process, both the value of the final product and the cost of producing the compound are important parameters dictating success (Scown *et al*, 2021; Ögmundarson *et al*, 2020). Currently only a narrow design space of compounds are economically feasible when purified sugars are used as the feedstock for bioconversion processes. As cells are genetically manipulated to produce new molecules, unanticipated “emergent” characteristics can appear as cells adapt to a new fitness landscape (Bajić *et al*, 2018; Wintermute & Silver, 2010; Rosenberg & Commichau, 2019; Guzmán *et al*, 2015), or in worst case scenarios, fail to produce the desired product.

To broaden the feasible product range to include bulk commodity chemicals, techno-economic analysis has indicated that lignin-based carbon streams may provide the cost savings to enable industrial scaleup and development (Corona *et al*, 2018; Bartling *et al*, 2021). Lignin valorization is a multifaceted problem. First, lignin is recalcitrant to depolymerization into individual carbon units that can be utilized in microbial bioconversion processes (Zhang *et al*, 2021; Cao *et al*, 2018). Next, the specific carbon units (furfurals, aldehydes, aromatics) resulting from lignin depolymerization are not consumed by many conventional microbes. Soil microbes like *Pseudomonas putida* KT2440 have emerged as excellent hosts for these processes due to their native catabolism of aromatics (and other non-glucose substrates) (Jiménez *et al*, 2002; Thompson *et al*, 2020). While engineered to produce a heterologous bioproduct, *P. putida* can utilize these non-conventional carbons streams to generate a wide range of molecules (Johnson *et al*, 2019; Almqvist *et al*, 2021; Loeschcke & Thies, 2015). The challenge addressed in this paper however, is how metabolic models can inform processes to increase productivity of final molecules from lignin-derived carbon streams. Metabolic models are mathematical representations of steady state metabolism and are historically informed using ^13^C-metabolic flux data obtained with positionally labeled isotopes of carbon substrates (Zamboni *et al*, 2009) and provide a generalizable and data-driven approach for strain engineering. The condition specific flux distribution requires information from the carbon molecule used for growth (i.e., glucose) which is not predictive of flux for other substrates (i.e., acetate or benzoate). Some relevant aromatic substrates, however, are prohibitively expensive and can cost up to $500,000 USD/gram (i.e., ^13^C-3-*para-*coumarate) when available. For aromatics, model accuracy has reasonable fidelity for products that can be derived from a few metabolic steps (Johnson *et al*, 2019) but for other products needing complex metabolic conversions (i.e. from a TCA node intermediate), confidence in model fidelity is low due to the absence of high-quality fluxomics data from aromatics alone. Some indirect methods have quantitated aromatic carbon flux using mixed carbon sources (Kukurugya *et al*, 2019), but they may not reflect metabolism when grown with aromatics as the sole carbon, which has received recent attention (Wilkes *et al*, 2023). Rigorously testing alternative methods for model refinement with lignin-derived carbon streams must be identified.

With regards to producing a specific desired molecule at high titer, rate, and yield, there are many sophisticated strategies that have demonstrated high product titers from a simple sugar (Meadows *et al*, 2016; Lo *et al*, 2016; Korman *et al*, 2017; Kim *et al*, 2019; Yim *et al*, 2011). The exact metrics necessary for process feasibility depends on a techno-economic analysis for the specific product (Yang *et al*, 2022). All three of these metrics are important for any scalable process, but few strain engineering techniques are able to address improvements to the rate. A low product rate is a concern for compounds that accumulate only in the stationary phase including bioplastics, dyes, and potential biofuel molecules (Zheng *et al*, 2013; Banerjee *et al*, 2020; García *et al*, 1999). We and others have demonstrated the use of computational strain optimization methods (Banerjee *et al*, 2020; Harder *et al*, 2016; Mehrer *et al*, 2019; Maia *et al*, 2016) to shift the mode of production from stationary phase to exponential phase, addressing this key concern. Our specific workflow utilizing growth coupled algorithms is called Product Substrate Pairing, or abbreviated PSP (Banerjee *et al*, 2020). While the principle of growth coupled methods call for complete inactivation of a potentially large set of reactions, we previously demonstrated that partial knockdowns or partial cut sets were sufficient to improve product rate formation by enabling production during growth phase (Banerjee *et al*, 2020). Here, we apply the PSP workflow for growth coupling followed by laboratory evolution, and systems-level high throughput -omics data analysis to optimize the bioconversion of a lignin-derived aromatic, *para-*coumarate (*p-*CA), to a commodity chemical and potential dye, indigoidine (Takahashi *et al*, 2007; Ghiffary *et al*, 2021). We achieved the growth coupled production of this molecule and exceeded 75% maximum theoretical yield from pure aromatics. Our engineered strains tolerate ionic liquid pretreated and base catalyzed depolymerized (BCD) sorghum liquor and produce indigoidine whereas the wildtype strain cannot.

## RESULTS

### Implementation of the PSP Workflow for the Bioconversion of *para-*Coumarate to Indigoidine

The results from this report are described in 3 major sections. First, computed growth coupled designs are prototyped using multiplex CRISPRi and down-selected for implementation as a deletion strain. Adaptive laboratory evolution was used to enhance growth of the generated triple deletion strain. In the second section, the best performing isolates were identified and their physical and cellular characteristics were enumerated. Finally, we use this characterization to directly improve final product titer, rate, and yield in an iterative strain engineering process.

### Selecting a Growth Coupled *p-*CA/Indigoidine Design for Implementation

In our previous study (Banerjee *et al*, 2020), we built a computational workflow (“Product Substrate Pairing,” or PSP) to assess if a new computational method for growth coupling (von Kamp & Klamt, 2017) was feasible in a soil microbe, *Pseudomonas putida* KT2440 using a genome scale metabolic model (GSMM), iJN1411. Our workflow assessed if the output cut sets were feasible for implementation in the lab and resolved metabolic reactions into specific genes, which in turn we could target for knockdown with multiplex CRISPR/dCpf1 interference (CRISPRi) in a multiplex format (Cpf1 is also referred to as Cas12a). Since publication of the earlier study, we made several improvements to the computational portion. First, we updated the genome scale metabolic model, iJN1411 to iJN1463, to include additional metabolic reactions. We also excluded duplicate reactions and updated the cMCS prediction algorithm to simply identify the genes instead of the enzymatic reactions. To account for *P. putida*’s native catabolism of *p-*CA to generate biomass as well as the heterologous indigoidine pathway, we customized the GSMM constraints. We allowed flux through the *p*-CA exchange and the intracellular reactions converting *p-*CA into 3-oxoadipate, which is routed to succinyl-CoA and acetyl-CoA as it enters the TCA cycle. We added the heterologous indigoidine production pathway to the GSMM and predicted the maximum indigoidine production potential in *P. putida* from *p-*CA and other major lignin derived aromatics. Using the GSMM iJN1463, maximum theoretical yield (MTY) of indigoidine was predicted to be 0.96 g/g of *p-*CA, 0.76 g/g of 4HBA, 0.85 g/g of ferulate, 0.74 g/g of vanillate and 0.82 g/g of vanillin (**Supplementary Table 1**), reflecting subtle differences in aromatic substrate catabolism.

To implement the PSP workflow, we assessed the solution space for a range of potential growth coupling solutions using both the constrained minimal cut set (cMCS) algorithm with the GSMM as well as elementary mode analysis (EMA) with the central metabolic network using a range of minimum biomass and yield constraints (**Figure 1A**, **Supplementary Figure 1**). The PSP workflow is agnostic to the specific computational optimization method and can evaluate predictions from any algorithm. Using the smaller network, EMA identified over 2,883 potential elementary modes but suffered from a low (<40% MTY) predicted theoretical indigoidine yield across a range of biomass formation rates. In contrast, the cMCS algorithm was computationally time-intensive, but out of 8 cMCS runs, amounting to over 1,231 cut set solutions, 810 of these had product yields exceeding 60% MTY (**Supplementary Figure 1**, **Supplementary Table 2**). Only two of these designed cut sets with >60% yields were down-selected based on our criteria in the PSP workflow (**Supplementary Table 3**) for being experimentally implementable (**Figure 1B**). In brief, we stipulated that the PSP designs selected would have fewer than 25 genes identified for intervention and at most 1 essential gene. After excluding the essential genes from the reaction sets, we built multiplex CRISPRi plasmids to test both of these designs and determined if gene knockdown would be sufficient to improve indigoidine production when cells were grown in M9 medium with *p*-CA as the sole carbon source. We tested these two plasmids and identified one of the Designs had up to a 4x improvement in indigoidine yield with subtle growth defects (**Supplementary Figure 2A, 2B, 2C**), but protein knockdown could not be validated by shotgun proteomics (**Supplementary Figure 2D**). Nonetheless, this prototyping information was sufficient to select one of these designs over the other for implementation as stable triple deletion strain and sidestep these protein quantification and detection limits encountered.

**Figure 1.**
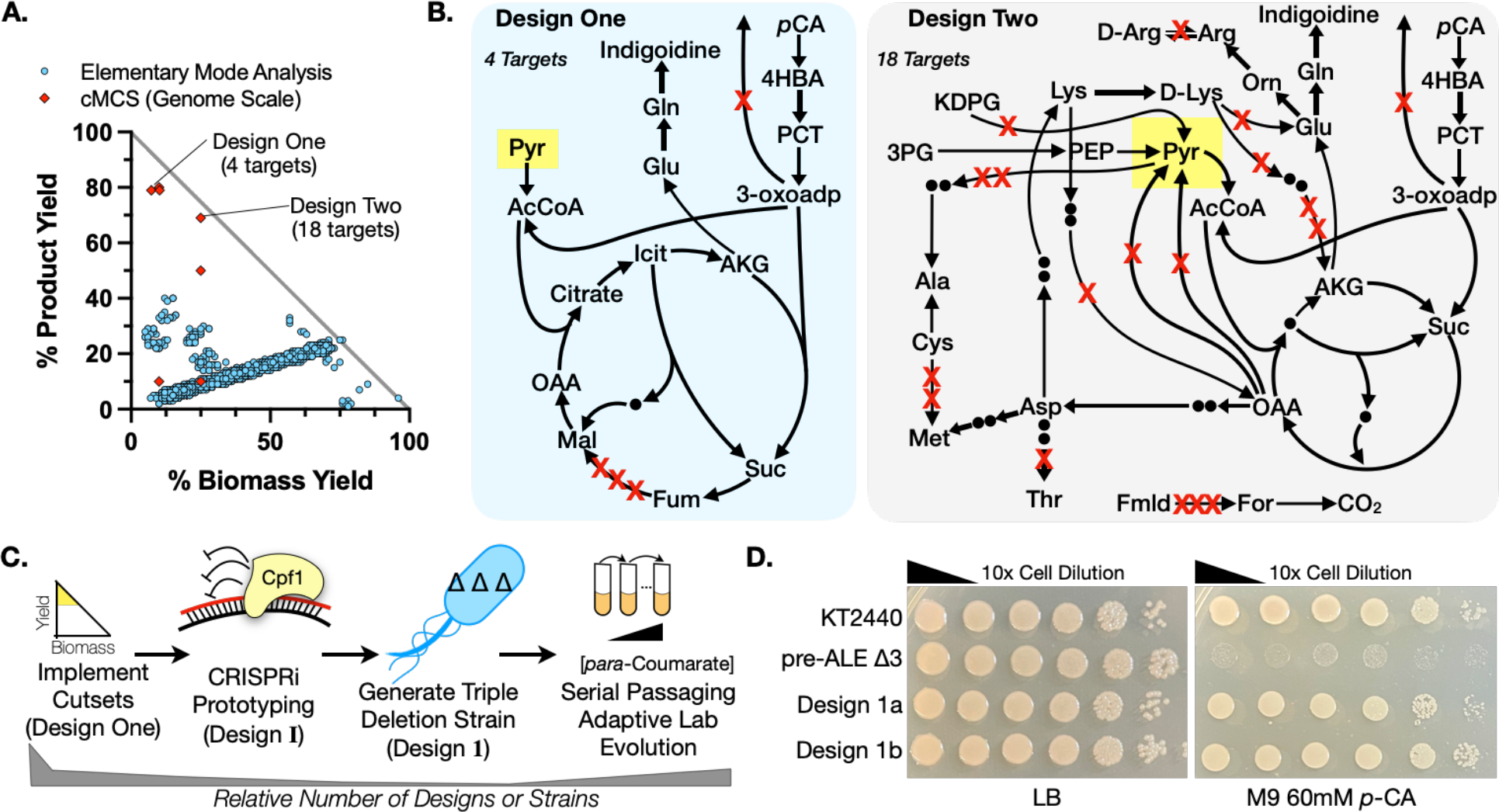
**Applying the PSP Workflow for the Growth Coupled Production of Indigoidine from *para*-Coumarate. (**A) Comparison of computational predictions using the constrained minimal cut set (cMCS) method vs. the elementary mode analysis (EMA) method. (B) Metabolic maps for PSP Designs One and Two selected for implementation. Gene targets are marked with red crosses. In Design One, three isozymes that catalyze the fumarate to malate reaction were targeted. (Refer Supplementary Table 3 for gene details.) (C) Diagram of workflow to compute, prototype, and select *P. putida* strains potentially growth coupled for indigoidine with *p*-CA. The height of the grey bar below indicates the relative number of strains or designs evaluated. (D) 10-fold serial dilution plating of *P. putida* pre-ALE triple deletion strain (pre-ALE Δ3) and two post-ALE triple deletion strain isolates (Design 1a and Design 1b) on LB and M9 60 mM *p-*CA solid agar media. WT *P. putida* (KT2440) is included as a control.

**Figure 2.**
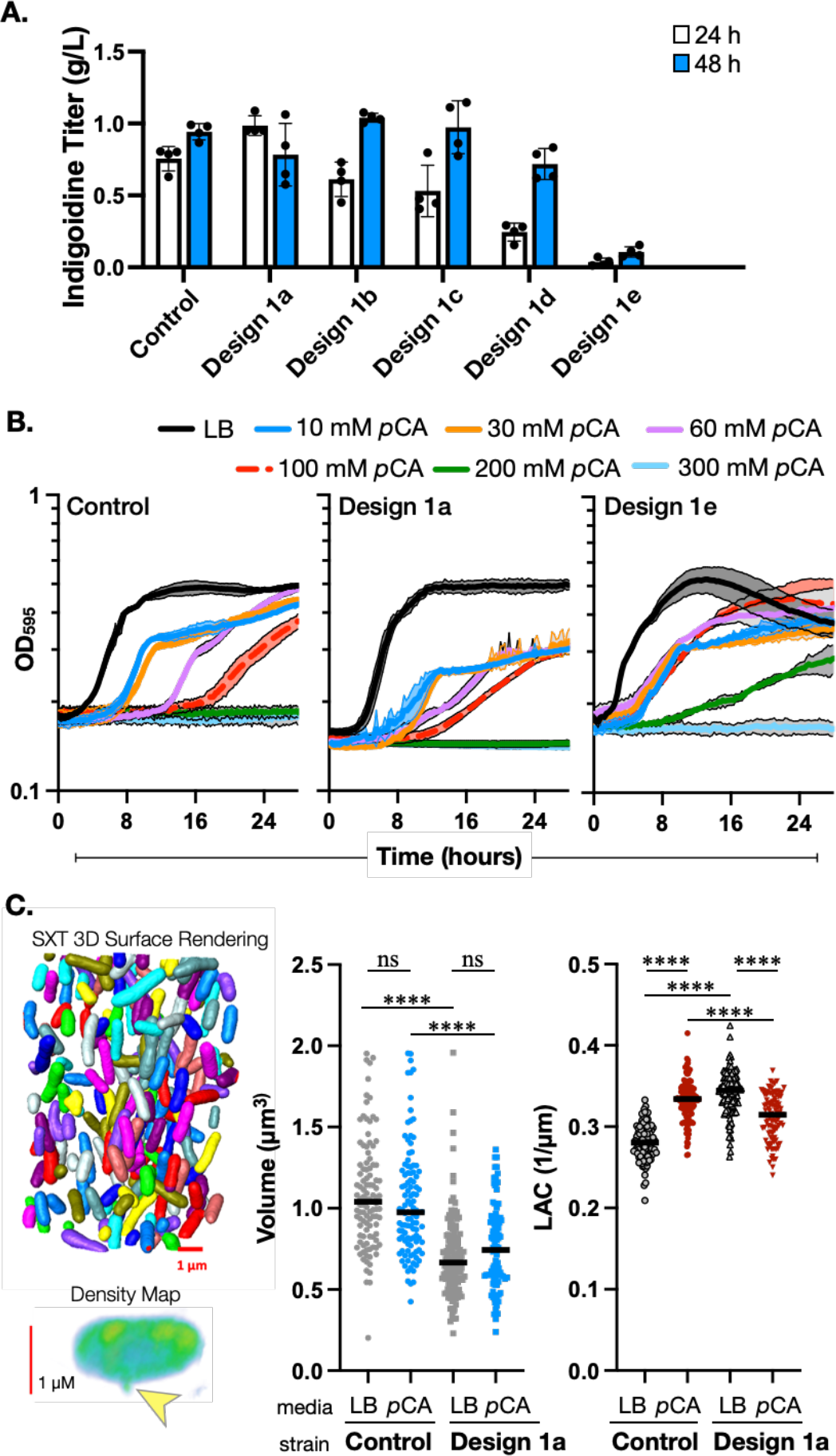
Initial Characterization and Down-Selection of Post-ALE Design 1 Isolates. (A) Indigoidine production was assayed across post-ALE Design 1 isolates (D1a, D1b, D1c, D1d, D1e) and compared to WT in a 24-well deep well plate incubated at 30 °C in M9 60 mM *p-*CA medium with 0.3% (w/v) arabinose. Indigoidine titer was quantified at the timepoints indicated. The error bar represents Mean ± S.D. (n = 4). (B) Post-ALE Design 1a and Design 1e strains were assayed for growth and tolerance at the *p-*CA concentrations indicated in M9 minimal medium in a microtiter plate. The shaded region between the color bands represents Mean ± S.D. (n = 3). (C) Soft X-ray tomography (SXT) analysis of Design 1a and WT strains in LB medium and M9 60 mM *p-*CA and 0.3% (w/v) arabinose. Left, top: 3D surface rendering from SXT. Left, bottom: A high density membrane aberration consistent in appearance with the formation of an outer membrane vesicle (OMV) is highlighted with a yellow arrow. Middle: Volumetric quantification. Right: Cell density analysis as measured by linear absorption coefficient (LAC). Strains and growth conditions for each sample are indicated on the bottom below the X- axis. The solid black line on the graph represents the average value. Ns, not significant. *P* < 0.0001 is represented by ****. At least 100 cells were examined for each dataset.

### Implementing Design One as a Triple Deletion Strain via CRISPR/Recombineering

Design One from the PSP pipeline identified a solution set of 4 genes for inactivation: 3 isozymes involved in the fumarate to malate node (PP_0897, PP_0944, and PP_1755), and a putative alpha-ketoglutarate /beta keto-adipate permease (*pcaT*/PP_1378). The predicted indigoidine yield from Design One is 76% MTY. PP_0897 is annotated as an essential gene and was excluded from this deletion set. Our base strain for engineering was the wild type *P. putida* KT2440 strain; the heterologous indigoidine pathway was integrated after all deletions were complete. We built this strain (ΔPP_1378 ΔPP_1755 ΔPP_0944; “pre-ALE Δ3” or “Δ3”) with CRISPR/Cpf1 aided recombineering (Materials and Methods, **Figure 1C**).The single mutant ΔPP_1378 strain grew slowly on M9 *p-*CA medium but had no discernible defects on LB (**Supplementary Figure 3A**). All possible double mutants combinations were viable but those including ΔPP_1378 were sick on M9 *p-*CA medium (**Supplementary Figure 3A**), while the triple mutant ΔPP_1378 ΔPP_1755 ΔPP_0944 strain failed to reliably grow on M9 *p-*CA plates, at most forming pinpoint or abortive colonies after six days incubation at 30 °C (**Figure 1D**). These results indicated that the triple mutant exhibited a synthetic lethality to *p-*CA compared to the double mutants.

Previously implemented growth coupled solutions can fail to grow in minimal medium as predicted, usually due to unexpected nutrient auxotrophies, which can be overcome by allowing the cells to generate spontaneous bypass mutants that overcome these growth defects (Harder *et al*, 2016; Luo & Hansen, 2018; Mehrer *et al*, 2019). We applied adaptive laboratory evolution (Sandberg *et al*, 2019) with 6 independent lineages derived from single colonies from several independently isolated clones in the Δ3 background as well as several backgrounds derived from this strain (see Materials and Methods and **Supplementary Figure 3B**). Since indigoidine can be visualized in cultures as a blue color, we were able to qualitatively monitor if indigoidine was still being produced after subsequent serial passages by including the inducer in the growth medium. We only recovered tolerant clones that still produced indigoidine in the Δ3 background strain (and in no other derivative backgrounds), and only when the cells were gradually shifted from LB medium without glucose to M9 *p-*CA medium. Curiously, several lineages could not be revived after cryostorage or failed to retain the improved growth in *p*-CA medium phenotype (**Supplementary Figure 3B**). These false positive clones were eliminated from further analysis.

### Characterization and Selection of Design 1 Strains by Multi-Modal Analysis

From our tolerization workflow we isolated spontaneous mutants with improved growth in M9 *p-* CA. These Δ3 isolates from ALE were assigned letter designations and will be referred to as Design 1x or abbreviated as D1x (the letters a-e indicate isolate designation). Strains D1a and D1b, the focus of this manuscript, were recovered after 18 or 19 serial passages respectively (**Figure 1D**), which is equivalent to approximately 90-95 doublings. Since these isolates were selected for growth under increasing concentrations of *p-*CA, we next screened mutants from the archival cryostocks to determine if the gain in *p-*CA tolerance modulated indigoidine production.

Indigoidine titers were measured in the deep well plate format where our CRISPR strains had shown high variability in product titers (refer to **Supplementary Figure 2A**) at the 24 and 48 hour timepoints post pathway induction (**Figure 2A**). The majority of our post-ALE strains showed comparable titers to WT with the exception of D1e, which was a poor producer (**Figure 2A**). Most strains produced around 1 g/L indigoidine under routine cultivation conditions, but D1e produced less than 100 mg/L indigoidine. We also examined tolerance of these isolates for growth at high concentrations of *p-*CA in M9 minimal medium (**Figure 2B**, **Supplementary Figure 4**). Strains D1a and D1b had comparable tolerance to WT, growing at 100 mM *p-*CA. The decrease in indigoidine titer correlated with this strain’s tolerance against high concentrations of *p-*CA. Wild type *P. putida* can grow in the presence of up to 100 mM *p-*CA, but isolate D1e exhibited robust growth at 200 mM *p-*CA and reduced lag times at concentrations equal to or lower than 100 mM *p-*CA. As there are many trivial reasons strains could have lost indigoidine productivity due to inactivating mutations in the heterologous pathway or inducer system, we selected the two best performing clones (D1a and D1b) for both highest absolute indigoidine titer and the best titer at 24 h, as it could reflect a kinetic shift towards growth coupled production.

### Outer Membrane Vesicles Are Detected, But Function Is Unclear

Recent work has indicated that *p-*CA may be catabolized in outer membrane vesicles (OMVs) that are secreted in response to an unknown lignin component (Alves *et al*, 2015; Vermaas *et al*, 2022; Xu *et al*, 2022; Salvachúa *et al*, 2020). We were curious if extracellular metabolite sharing was playing a role in our Design 1 strains with restored growth on *p*-CA. We tried to detect OMVs and other structural changes by means of high-resolution imaging using soft X-ray tomography (SXT). SXT generates 3D structural information of intact cells at mesoscale resolution (∼25 nm) using cryo-fixed cells without any fixatives as reviewed previously (Loconte *et al*, 2023). In addition to structural data, SXT provides quantitative information about the density and composition of organic molecules. From this method we observed structural and morphological differences in composition compared to the Control strain as well as the appearance of dense membrane structures consistent with previous reports of OMVs in the process of budding (**Figure 2C**, **Supplementary Figure 5A**). We were able to observe membrane-bound buds approximately 270 nm in size (**Supplementary Figure 5A**) emerging from both WT and Strain D1a, matching the reported OMV dimensions (Salvachúa *et al*, 2020). In both LB and M9 *p-*CA media, cells from D1a were volumetrically smaller and had increased density when quantified with the linear absorption coefficient (LAC) (**Figure 2C**). Strain D1a showed decreased cell density in M9 *p-*CA medium compared to LB medium. In contrast, the control strain showed increased density from LB to M9 *p*-CA medium. While Design 1a cells are smaller volumetrically, there was no change in colony forming units (CFUs) when quantified by serial dilution plating (**Supplementary Figure 5A**). Changes in cell density have been previously reported correlating with cell cycle progression in mitotically synchronized fission yeast cells, but the biological relevance related to restoring *p-* CA catabolism is not clear (Odermatt *et al*, 2021). From this physical characterization we conclude that the changes in *p*-CA tolerance and the resulting restoration of indigoidine production have generated persistent changes in cell shape and density that are apparent under rich medium cultivation conditions, and that nascent OMVs can be detected in cells frozen in their near-native states without crosslinking reagents.

Diffusible molecules that are exchanged between adjacent strains on an agar plate can be detected if they change the growth patterns of the proximal microbe (Eng *et al*, 2020). We tested the metabolite exchange hypothesis using a Δ*fcs* strain (deficient in the first step of *p-*CA catabolism) to measure if this Δ*fcs* strain could access *p*-CA derived metabolites generated from adjacent WT on an agar plate. Our observations indicate that extracellular metabolic processes potentially mediated by OMVs are not reliably functional at biologically relevant timescales in either WT or strain D1b when grown in this interaction assay (**Supplementary Figure 5B, 5C**), and is consistent with a similar liquid culture report measuring OD changes from Salvachua *et al*.

### Hybrid Whole Genome Sequencing Analysis

The pre-ALE Δ3 strain displays a synthetic lethality to *p-*CA as the carbon source as the single mutants and double mutants are viable, and with ALE we were able to restore growth. There were two aspects of our strains that required characterization: first, what were the mutations that restored tolerance to *p-*CA as a carbon source and secondly, how did cell metabolism rewire itself in response to compromising 2 nodes in central metabolism? To characterize the genetic changes in these strains and their downstream impact on protein expression, we applied both whole genome resequencing and shotgun proteomics to capture changes in protein function (via non-synonymous codon changes) as well as protein abundance as a proxy for increased activity. For DNA sequencing, we used hybrid Nanopore long read sequencing in conjunction with Illumina short read sequencing for *de novo* genome assembly and annotation (**Figure 3A** and Materials and Methods). High quality contigs were assembled (Materials and Methods, **Supplementary Figure 3C**) and polymorphisms were analyzed for correlations with restored growth on *p-*CA (**Supplementary Results**).

**Figure 3.**
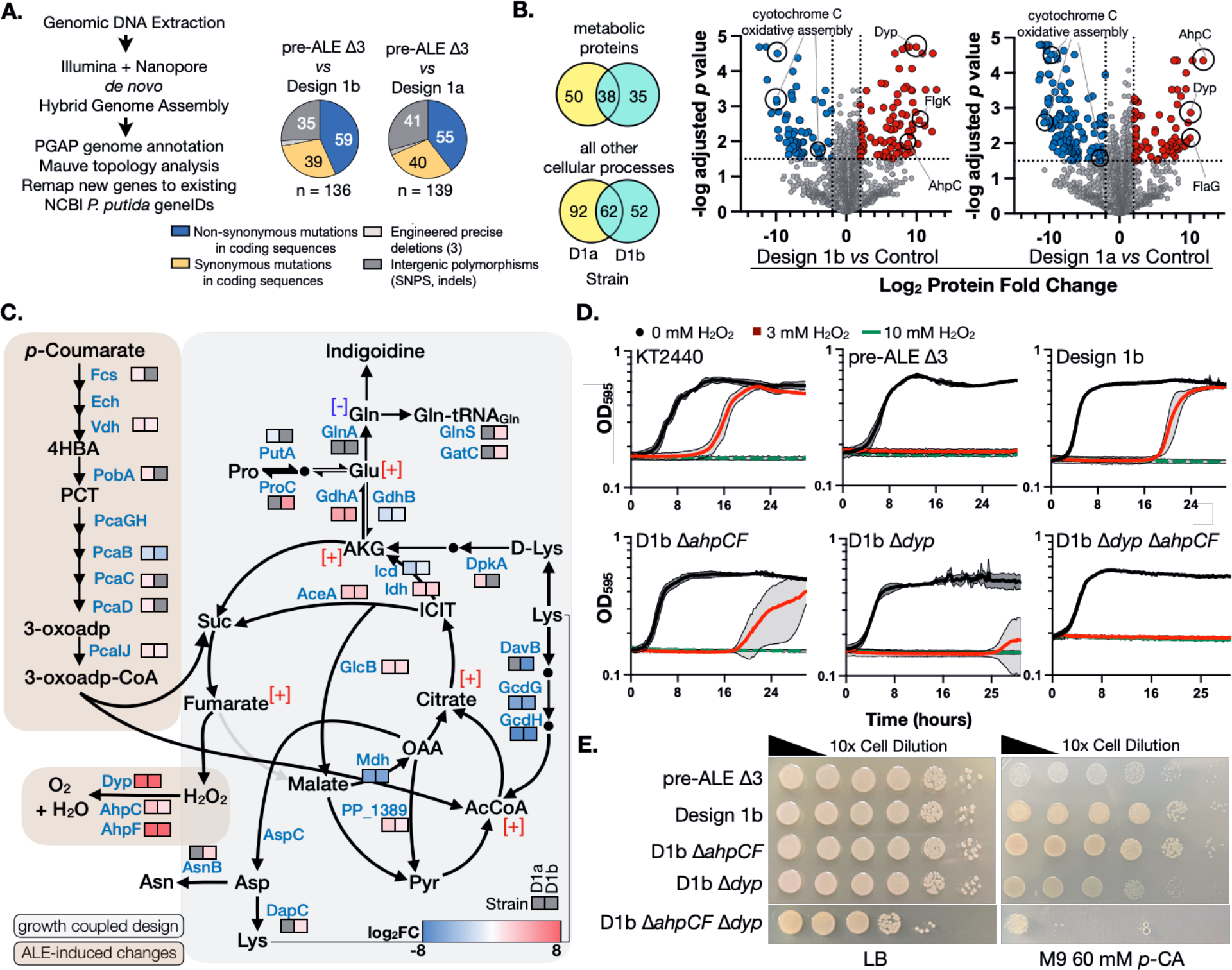
Multi-Omics Analysis of PSP Design 1 Clones a and b. (A) Left. Hybrid genome assembly and annotation workflow. Middle. Pie charts comparing pre-existing mutations in the pre- ALE Δ3 strain to the two post-ALE isolates, Design 1a (D1a) and Design 1b (D1b) clones. (B) Left. Venn diagram of significantly enriched proteins for D1a and D1b. Right. ∼2,200 proteins detected by LC-MS/MS of D1b and D1a compared to the Control strain. (C) Proteomics analysis of inferred changes in central metabolism due to ALE (brown box) or Design 1 based gene deletions (gray box). Key metabolites in a D1b derivative strain were quantified by LC-MS and metabolite concentrations higher than 2-fold are indicated with [+] in red; and reduced by 5-fold is indicated with [-] in blue. (Refer Supplementary Figure 7B for double deletion strains). (D) Top. Peroxide sensitivity of WT *P. putida*, pre-ALE Δ3, and D1b isolates to exogenous hydrogen peroxide (H2O2) in a kinetic growth assay. Concentrations of hydrogen peroxide in LB medium were 0 mM, 3 mM and 10 mM. Bottom. Single and double deletion mutants lacking peroxidase encoding genes *ahpCF* and/or *dyp* in the D1b background were also tested. The shaded region between the solid line indicates Mean ± S.D. (n = 3). (E) 10-fold serial dilution of *P. putida* peroxidase mutant strains in D1b background on LB or M9 60 mM *p-*CA plates. The pre-ALE Δ3 strain is included for comparison.

There are 3 specific mutations in these post-ALE Design 1 strains that we draw attention to. First, we identified point mutations in the upstream promoter sequence of PP_1378, which was previously targeted for deletion in the Design 1 strains. Strain D1a contained a G/A substitution at the -6 position upstream of the PP_1378 promoter, while strain D1b contained a C/G substitution at the -12 position upstream of the same gene. In the process of removing PP_1378, we may have impacted the expression of the remaining genes in the PP_1378-PP_1381 (*pcaTBDC*) operon even though we left the promoter sequences upstream of the PP_1378 start codon intact. This operon is crucial to the conversion of intermediate metabolites 4-hydroxybenzoate (4-HBA) to protocatechuate (PCT) in the *p-*CA catabolism pathway. The ΔPP_1378 single mutant strain grows poorly on M9 *p-*CA medium (**Supplementary Figure 3A**), consistent with the need to accumulate an activating mutation in this *pcaTBDC* promoter region. Second, we detect gain of function mutations in two transcriptional regulators PP_3493 and PP_4338. RB-TnSeq analysis indicates their inactivation leads to weak fitness defects on different aromatics but not glucose (**Supplementary Figure 6A**); it is possible the mutants here increase their activity. The PP_3493- N215S mutation impacts the protein substrate recognition domain while PP_4338-N72S impacts the phosphotransferase domain of a chemotaxis-related histidine kinase, which could modulate the activity of this signaling cascade. *P. putida* is known to induce chemotaxis towards aromatic molecules with some cross-activation of their cognate catabolic pathways (Luu *et al*, 2015). These three mutations, one in *p-*CA catabolism and two transcriptional regulators, are the most likely genes implicated in restoring growth on *p*-CA in these two isolates by DNA resequencing.

### Emergent Properties of the Design 1 Strains Identified Using Shotgun Proteomics

The key finding from the proteomics analysis was the identification of a new route for how carbon was rewired from *p*-CA to indigoidine. The fumarate to malate node was bypassed by increasing proteins needed for the glyoxylate shunt pathway, redirecting flux from lysine towards AKG, and in turn favoring glutamine and indigoidine formation.

Shotgun proteomics revealed 123 metabolic and 206 non-metabolic proteins with differential protein counts (**Figure 3B**). While there were clear correlations of upregulated aromatic catabolic pathway proteins known to have fitness defects across all aromatic molecules profiled by RB-TnSeq (**Supplementary Figure 6B**), there was no correlation between the differential protein changes for TCA cycle, glycolytic, homogentisate or metal cofactor proteins profiled.

To assess changes in the initial *p-*CA catabolism, we compared all possible double mutant pairs with the Design 1b strain using proteomics. Both double deletion strains lacking PP_1378 had reduced counts (between 2-8 fold log2) for the proteins responsible for converting 4-HBA to PCT in *p-*CA catabolism (**Supplementary Figure 7A, 7B**). This is consistent with their slow growth phenotype on M9 *p*-CA medium (**Supplementary Figure 3A**). In contrast the ΔPP_1755 ΔPP_0944 strain expressed higher levels of PcaB than strain D1b, with no growth defect on M9 *p-*CA. The restored growth and protein levels in the Design 1 strains correlates with the unique point mutations recovered in the promoter sequence upstream of PP_1378 described for strains D1a and D1b. The decrease in pathway protein abundance for several other steps in the conversion of 4-HBA to PCT suggests the loss of *pTBDC* operon expression impacted other pathway proteins including PobA, PcaIJ, and PcaGH. We conclude that the Design 1 strains restored conversion of *p-*CA into 3-oxoadipyl-CoA as a consequence of ALE.

Proteomics also suggested there was an active shunt of the carbon flux towards succinate and malate to bypass the inactive part of TCA due to the targeted gene deletions (**Figure 3C**).

Examining the Design 1a and 1b strains *vs.* WT control dataset, we observed differences in the conversion of fumarate to malate. PP_0944 activity was reduced below the detection limit consistent with its expected gene deletion. PP_0897, the third fumarate to malate isozyme, had 25% - 30% higher protein counts in both D1 isolates than WT. At the malate to oxaloacetate node, malate dehydrogenase, (Mdh/PP_0654) had zero protein counts; Mqo3 protein had a significant fold reduction (only 9% to 26% that of WT) whereas the other two quinone dependent malate oxidoreductase proteins, Mqo1 and Mqo2, were slightly elevated in strains D1a and D1b compared to WT. This implies that malate is rerouted into pyruvate (MaeB/PP_5085, nearly 2-fold higher protein abundance). 3-oxo-adipyl-CoA can also enter the TCA cycle through acetyl-CoA catalyzed by GltA. We detected increased protein counts in both D1 isolates strains compared to WT. Citrate to isocitrate proteins AcnA1, AcnA2, AcnB, had similar or higher protein counts in both D1 isolates compared to WT with the exception of AcnA1 in strain D1a (reduced to 10% of WT). Isocitrate to alpha-ketoglutarate (AKG) metabolic step involves the isocitrate dehydrogenase isoforms Idh/PP_4012 and Icd/PP_4011. Idh was upregulated while Icd was downregulated. AlphaFill analysis (Hekkelman *et al*, 2023) suggests that the differences in protein enrichment might be due to differential cofactor specificity for either NADH or NADPH (**Supplementary Data 1**). Isocitrate through the glyoxylate shunt via protein AceA/PP_4116, was upregulated (>2.59) and GlcB (>1.57), also had 3-fold higher protein counts. Proteins involved in AKG to succinyl-CoA (SucA, SucB and LpdG) were reduced by 20 - 50% of protein counts compared to WT. Overall, all the proteins involved in TCA were similar or higher than WT protein counts except for the expected deleted genes and Mdh/PP_0654.

Finally, several proteins related to the downstream metabolite pools needed for indigoidine formation were upregulated in the rewired strain. Indigoidine is derived from the condensation of two glutamine molecules, which are generated from glutamate and in turn AKG. Glutamate dehydrogenase GdhA/PP_0675 (>4.61), was highly upregulated in both D1 isolates while GdhB/PP_2080 (<-1.42) was downregulated. Glutamine synthetases, GlnA and PP_4399, had decreased protein counts to 50% - 60% in D1a and 8% - 30% in D1b of WT levels. Lysine re-entry into the TCA cycle via the AKG node was favored over acetyl-CoA. DpkA/PP_3591 was highly enriched in the Design 1 strains, with raw protein counts higher than 2.79-fold compared to WT whereas lysine metabolism towards acetyl-CoA via glutarate metabolic proteins DavB/PP_0383, GcdH/PP_0158 and GcdG/PP_0159 was significantly reduced or undetected in both the D1 isolates. Glutamate metabolism towards glutathione biosynthesis through protein GshA was four-fold higher than the WT control. Levels of glutathione are important in maintaining redox balance and are closely linked to oxidative stress responses in *P. putida* (Nikel *et al*, 2021). These changes in lysine metabolism are unexpected and indicate how the unintuitive deletions suggested from the computational predictions have implications for other aspects of cell physiology and metabolism.

The relative metabolite concentrations agreed with the implied changes from the proteomics analysis (**Supplementary Data 2**). Reduced flux through the fumarate to malate node was confirmed with a 2.7-fold increase in fumarate concentration in strain D1b compared to WT. Acetyl-CoA and citrate had increased by 2-fold, while AKG had increased by 2.71-fold. Finally, glutamate had increased by 4.4-fold and glutamine decreased by 6-fold. This D1b derivative strain expresses an additional copy of *glnA*. These metabolite concentrations provide important evidence corroborating the differential protein expression as a proxy for metabolism we have observed in these Design 1 strains.

### Upregulated Peroxidases Are Necessary to Remedy Oxidative Stress in Design 1 Strains

The class of upregulated oxidative stress proteins suggest a mechanism for how growth was restored in the Design 1 strains. Two types of peroxidases (Dyp/PP_3248; AhpCF/PP_2439, PP_2440) were highly upregulated in both Strains D1a and b (**Figure 3B**, **Supplementary Figure 6B)**. Peroxidases catalyze the reduction of hydrogen peroxide to water and oxygen and for organic hydroperoxides to water and alcohols (Chubukov *et al*, 2015). Our strains accumulate fumarate due to the deletion of two key enzymes that convert fumarate to malate (**Figure 3C**). Fumarate reductase is a major source of hydrogen peroxide in *E. coli*, and the *E. coli* AhpC homolog is known to ameliorate these peroxides during aerobic growth (Forrester *et al*, 2018; Esterházy *et al*, 2008; Meehan & Malamy, 2012; Messner & Imlay, 2002; Korshunov & Imlay, 2010). We hypothesized that growth on *p-*CA medium generates additional reactive oxygen species normally remedied as part of the electron transport chain reaction; we tested this hypothesis by interrogating if these pre and post ALE strains were differentially sensitive to hydrogen peroxide. Moreover, deleting the peroxidases would resensitize the strains to peroxides as well as block growth with *p-*CA as the sole carbon source.

Using an exogenous peroxide stress assay (Materials and Methods), we compared the ability of our strains to grow in the presence of this environmental insult. WT *P. putida* strains were able to grow in the presence of 3 mM hydrogen peroxide, but our pre-ALE Δ3 strain was sensitive to this environmental stressor and failed to grow (**Figure 3D**). As controls, the single mutants ΔPP_1755 and ΔPP_1378 or the ΔPP_1755 ΔPP_1378 double mutant strain were not sensitive to 3 mM hydrogen peroxide (**Supplementary Figure 7A**). Consistent with this hypothesis, growth was restored with 3mM hydrogen peroxide in the Design 1b strain. Deleting either of the peroxidases (*ahpCF* or *dyp*) in the Design 1b strain impaired growth in peroxide-containing media; deleting both peroxidases in the D1b strain reduced growth to the same background level as the pre-ALE strain. The D1b *ΔahpCF* strain did not show any growth defect on M9 *p*-CA plates but D1b Δ*dyp* strain showed a pronounced growth defect that formed pinpoint colonies after three days (**Figure 3E**). The double mutant strain, D1b Δ*ahpCF* Δ*dyp*, failed to grow on M9 *p-*CA medium, only forming colonies on the first spotted dilution with no background growth at higher dilutions, confirming our hypothesis. These results indicate that adaptive laboratory evolution is needed to overcome additional reactive oxygen species generated as a consequence of the growth coupled design and that upregulating both peroxidases was necessary to restore growth to wildtype levels.

Due to the growth defect in the D1b Δ*dyp* Δ*ahpCF* strain, we wondered if peroxidase activity was sufficient to restore growth in the Δ3 strain without undergoing ALE. When *dyp* was overexpressed in the Design 1b Δ*dyp* strain, growth on M9 *p-*CA was restored compared to the empty vector control (**Supplementary Figure 7B**). However, when the same construct was expressed in the pre-ALE Δ3 strain, the cells grew worse than the empty vector control. It was unlikely that over- expressing the peroxidase would be sufficient to restore growth, since we know there are many other compensatory mutations in the Design 1 strains related to the uptake and initial catabolism of *p-*CA (ie, PP_4338-N72S aromatic molecule sensing or PP_1378 promoter mutations related to the *pcaTBDC* operon). We conclude the peroxidases are necessary but insufficient for restoring growth in the Design 1b isolate.

### Increasing Growth Coupled Indigoidine Production with Rational Engineering Methods

We next interrogated the relationship between growth and indigoidine final product yield in our Design 1 strains. The basic parameters for cultivation and product characterization were established by generating a new indigoidine standard curve produced from wildtype *P. putida* KT2440 expressing the heterologous indigoidine pathway (“Control” strain) when fed *p-*CA (see Materials and Methods and **Supplementary Figure 8A**, **8B**). Cultivation medium parameters were also evaluated; a range of starting *p-*CA concentrations of 60 mM to 120 mM *p-*CA was optimal when either ammonium sulfate or ammonium chloride was used as the nitrogen source, and 100 mM ammonium yielded slightly improved titers of 1 g/L indigoidine with tighter confidence intervals (**Supplementary Figure 8C**). The major difference in culture media between the initial production data shown in Figure 2 with the optimized condition was a switch from 15 mM ammonium sulfate to 100 mM ammonium chloride. We also developed an orthogonal LC-MS based method to quantify indigoidine based on its mass and fragmentation pattern (**Supplementary Figure 8D**). Purified indigoidine does not contain any detected contaminants as the quantification from the colorimetric assay strongly correlates with the reported values from LC-MS analysis (R^2^ = 0.9984) (**Supplementary Figure 8D**).

The desired outcome for any growth coupled strategy is shifting the production phase to the exponential growth phase. To interrogate if indigoidine became growth coupled when fed *p-*CA in the Design 1 strains, we prepared samples for a timecourse analysis to characterize production during exponential growth. Indigoidine yield from both Design 1 strains were compared to the control *P. putida* production strain. While Strain D1a and D1b produced indigoidine at the 8 hours timepoint (7 mg indigoidine/g *p*-CA and 75 mg indigoidine/*p-*CA, respectively), WT control strain did not (**Figure 4A**, left panels). We confirmed that at the 5 and 8 hours timepoints all three strains were still consuming *p-*CA until the 24 hours timepoint (**Figure 4A**, right panel). We note that the control strain at this same mid-log timepoint had 5-fold more Sfp indigoidine production pathway protein expression than Design 1b, indicating that production pathway protein expression alone is insufficient to drive indigoidine formation during the growth phase (**Supplementary Data 2**). The shifting of indigoidine from stationary phase to log phase production extends our initial observation from glucose (Banerjee *et al*, 2020) to an aromatic carbon stream.

**Figure 4.**
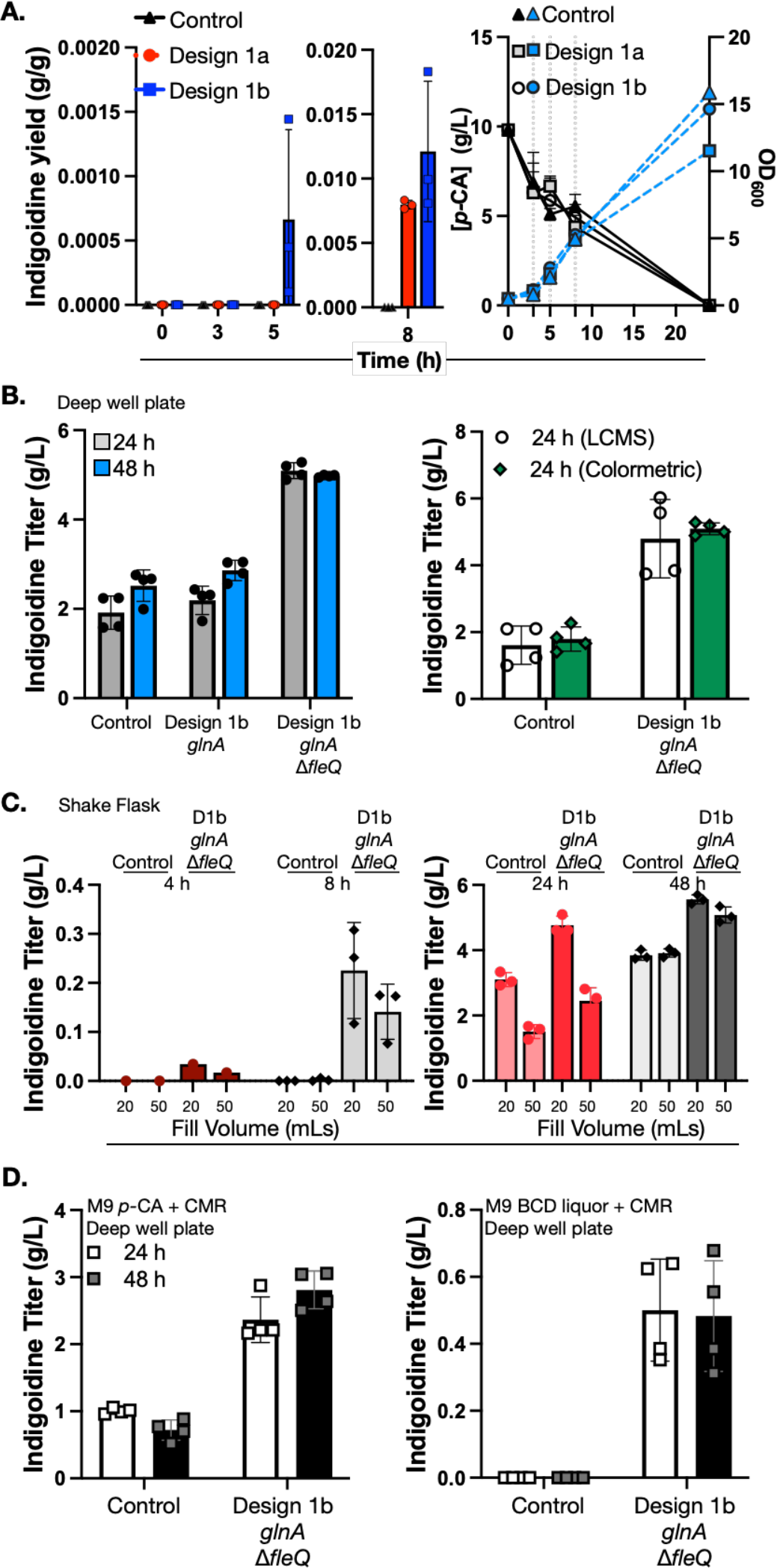
Demonstration of Growth Coupled Bioconversion of *p-*CA to Indigoidine and Rational Strain Engineering Approaches to Increase Titers. (A) Indigoidine production kinetics in shake flasks with 0.3% w/v arabinose. The indigoidine yield was calculated as grams per gram (g/g) of *p-*CA consumed at the given timepoint. (B) Left: Indigoidine assay for Design 1b augmented with *P. stutzeri glnA*, and deletion of *fleQ,* the master regulator of the flagella biosystem. Arabinose inducer concentration was increased from 0.3% to 1.5% w/v. Right: Titers from the colorimetric method were cross-validated using LC-MS. (C) Indigoidine production kinetics of Design 1b *glnA* Δ*fleQ* compared to WT, as in (A) at the timepoints indicated in 20 mL and 50 mL fill volumes in 250 mL shake flasks. Arabinose was used at 1.5% w/v. (D) Indigoidine production for *P. putida* Control strain harboring the indigoidine production pathway and the Design 1b *glnA* Δ*fleQ* strain using base catalyzed depolymerization (BCD) sorghum liquor. Arabinose was used at 1.5% w/v. (Refer Supplementary Table 5 for aromatic carbon composition of BCD liquor). Error bars represent Mean ± S.D. (n=3) in A and C and Mean ± S.D. (n=4) in B and D.

While our Design 1 strains here improved the overall rate, they did not meaningfully increase the final product titer (refer to **Figure 2**). We first addressed this by taking a rational strain engineering approach by examining precursor supply, as glutamine synthetase protein counts were somewhat reduced in the Design 1 strains (**Figure 3C**). We augmented the indigoidine production pathway by introducing the glutamine synthetase *glnA* from either *P. stutzeri* RCH2 or *P. putida* KT2440 downstream of the existing pathway to form a 3-gene operon under the control of the BAD promoter. The introduction of *glnA* from *P. stutzeri* RCH2 had the most robust improvement to indigoidine titer in both Design 1 strains, increasing titers up to 2-fold to 2 g/L and therefore chosen for following strain engineering (**Supplementary Figure 9A**). The same three gene indigoidine pathway design had limited impact on titer when expressed in the WT control *P. putida* (**Supplementary Figure 9B**), suggesting GlnA activity was only rate limiting in the Design 1 isolates consistent with our proteomics analysis. Next, we varied the arabinose inducer concentration, and observed that higher indigoidine titers were possible by increasing the concentration from 0.3% to 1.5% arabinose, but here we only observed an improvement in the control strain and not the Design 1b strain (**Supplementary Figure 9B**).

The proteomics dataset provided additional targets for greater improvements to indigoidine titer. Both Design 1 strains showed many flagella related proteins were upregulated log2 4-10 fold over the WT control. In WT *P. putida*, approximately 8 percent of energy in a cell is devoted towards cell motility and is controlled by the FleQ transcriptional master regulator (Blanco-Romero *et al*, 2018; Martínez-García *et al*, 2014) . We tested indigoidine production in our Design 1b *glnA* strain after deleting *fleQ* and found that titers increased from 2 g/L to ∼5 g/L in this new strain (**Figure 4B**). The titer values from the deep well plate were corroborated by our LC-MS method. The control KT2440 Δ*fleQ* strain in comparison only showed a 30% indigoidine titer improvement over the control, likely reflecting the increased flagella-based metabolic burden in the Design 1 strains (**Figure 3B**, **Supplementary Figure 9B**). The D1b *glnA ΔfleQ* strain showed stable indigoidine titers across cultivation formats in both the deep well plate and shake flask formats with similar titers at 5.1 g/L at the 48 hours timepoint (**Figure 4B, 4C**). In shake flasks, a decreased total fill volume led to higher early titers (200 mg/L) compared to the original D1b strain at the 8 hours timepoint (**Figure 4A**). The strains used in **Figure 4C** in the 20 mL shake flask setup were analyzed for metabolite concentrations that were described earlier in **Figure 3C**. These results indicate the robustness of *p-*CA to indigoidine bioconversion in the Design 1b *glnA ΔfleQ* strain under different cultivation formats.

Finally, we used strain D1b *glnA ΔfleQ* to produce indigoidine from real-world lignin derived carbon streams by replacing commercially purchased *p-*CA with sorghum biomass-generated BCD liquor (Materials and Methods). The concentration of *p-*CA present in 1x M9 BCD liquor medium was expected to be around 138 mg/L, which is suboptimal for indigoidine production based on our previous work in C/N ratio (**Supplementary Figure 8C**). Furthermore, there are many other characterized aromatic molecules present in BCD liquor at similar concentrations that might also be co-catabolized with *p-*CA (**Supplementary Table 5**), which may or may not be compatible carbon substrates for the initial growth coupled design (**Supplementary Table 4**). While both the control strain and the D1b *glnA ΔfleQ* strain were able to grow in M9 BCD liquor (**Supplementary Figure 10**), strain D1b *glnA ΔfleQ* was able to reach somewhat higher densities as assayed by serial dilution and semi-quantitative CFUs at the 24 h timepoint by serial dilution of cultures onto LB agar medium (**Supplementary Figure 9C**). Importantly, we observed in this batch of M9 BCD liquor, only the D1b *glnA ΔfleQ* strain was able to produce 500 mg/L indigoidine within 24 hours, while WT did not produce detectable indigoidine even after 48 hours (**Figure 4D**). This result suggests the D1b *glnA ΔfleQ* strain was more tolerant of BCD liquor containing a mixture of carbon streams for indigoidine production, whereas WT failed to produce any indigoidine with this batch of sorghum BCD liquor. As this yield exceeds the initial *p-*CA concentration in M9 BCD liquor, we presume that other carbon sources present in BCD liquor were used to generate this quantity of indigoidine.

### Refining Genome-Scale Metabolic Model Constraints with Proteomics

To fully utilize all information generated from the proteomics data, we adapted an existing workflow for integrating transcriptomics or proteomics data into genome scale metabolic models. Using the iMAT algorithm (Zur *et al*, 2010) we integrated the GSMM with proteomics data which generated a flux activity state for each reaction in the model, reconstraining the presence or absence of the metabolic flux annotated in the GSMM for the wild type strain. We generated two context-specific models representing strain D1a (iMATD1a559) and strain D1b (iMATD1b539) (**Figure 5A**). The resulting iMATD1b539 model reduces the active 1644 reactions in the original GSMM under *p-*CA minimal medium condition, to 558 reactions. This model contains 558 reactions, 543 metabolites and 539 associated genes. We also considered using proteomics data generated from derivative strain D1b *glnA* Δ*fleQ,* but this strain showed very few non-metabolic proteins with differential protein expression levels compared to the parental D1b *glnA* strain, indicating this flagellar master regulator does not regulate many metabolic processes (**Supplementary Data 2**). Satisfied that the Δ*fleQ* deletion would not change the differential protein counts used for constraints, we used the iMATD1b539 model to compute a second PSP cycle for growth coupled production of indigoidine from *p*-CA. We computed cut sets using a minimum of 80% MTY of indigoidine and a maximum of 25% MTY biomass yield. This resulted in the smallest cut set with a total of 4 metabolic genes: PP_4120, PP_0158, PP_0597 and PP_0253. PP_0253 was excluded as it is a pseudogene, and we note that the PP_4120 deletion was previously identified as a single deletion that can both improve indigoidine titer and fitness in a bioreactor (Eng *et al*, 2021). As before we sequentially generated the next three deletions in the D1b *glnA* Δ*fleQ* strain and tested the resulting deletion strains for indigoidine production. While the new strains containing one additional deletion did not show appreciable titer increases, strain D1b *glnA* Δ*fleQ* PP_4120 ΔPP_0158 increased titers from 4.5 g/L indigoidine to 6.2 g/L, equivalent to 65.6% MTY (**Figure 5B**). The completed cut set strain D1b *glnA* Δ*fleQ Δ*PP_4120 ΔPP_0158 ΔPP_0597 increased indigoidine titers even further to 7.3 g/L, equivalent to 77% of the MTY from *p-*CA (**Figure 5B**). Unlike the previous Design 1b isolates from earlier iterations, we now observe a growth defect in this strain with smaller colonies and fewer absolute number of CFUs on M9 *p-*CA plates as well as slower growth in liquid cultures (**Figure 5B**, **Supplementary Figure 9D**), suggesting a tradeoff between high indigoidine titers and biomass formation.

**Figure 5.**
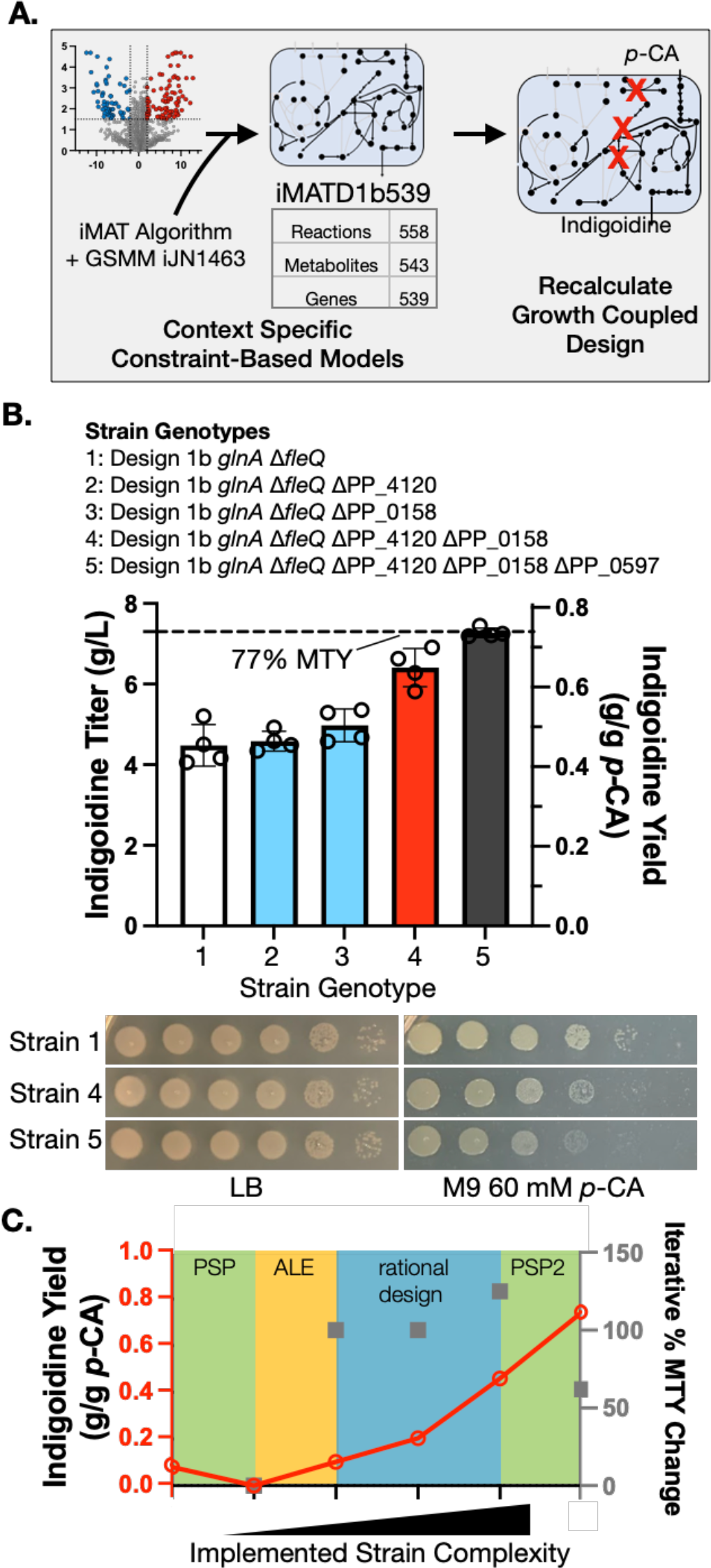
Context-specific Models Using Proteomics Constraints Led to Maximal Indigoidine Titers. (A) Model-based proteomics data integration pipeline. In a second PSP cycle (PSP2), we recomputed gene deletion targets for the growth coupled production of indigoidine from *para*- coumarate using a new proteomics-informed, context-specific metabolic model (iMATD1b539). (B) Indigoidine production profile (top) and 10-fold serial dilution plating assay (bottom) for *P. putida* strains generated from the second cycle of PSP. Samples were grown in deep well plates and analyzed for indigoidine production from the 24 h timepoint. 1.5% w/v arabinose was used as the inducer. The strain genotypes assayed are indicated in the Figure. The error bars represent Mean ± S.D. (n=4). (C) Summary of cumulative indigoidine yield gains (solid red line) and iterative % MTY change for each subsequent strain generated (grey squares, (**▪**)). The color blocks indicate the method of strain engineering used. PSP: best multiplex CRISPRi prototype and initial triple deletion strain. ALE: post ALE Design 1 isolates, highest titer. Rational Design: C/N ratio optimization, integration of *P. stutzeri glnA*, & deletion of *fleQ.* PSP2: Three additional deletions based on the second cycle of PSP (PSP2) using iMATD1b539.

## DISCUSSION

Genome scale metabolic models are comprehensive representations of cellular metabolism that have potential for use in the predictive design of microbes. GSMMs have been extended to heterologous pathways and many other organisms (Maia *et al*, 2016; Banerjee & Mukhopadhyay, 2023). These methods are ideal for optimizing bioconversion processes used to produce commodity chemicals as most require high productivity using a range of carbon sources. Designing microbes for this purpose is nontrivial and the execution is rarely predictable.

Here we implemented growth coupling for both the expression of a heterologous product and the use of a non-canonical carbon source. From our previous work (Eng *et al*, 2021) we anticipated that many important non-metabolic processes would impact final product titers but were unaccounted for in the genome scale metabolic model. We affirm that the current state of the *P. putida* GSMM and tools for growth coupling are well poised for effective strain engineering. Applications where metabolic flux information (and model confidence) is limited can be remedied with these methods to produce final products distal from their starting substrates. The initial objective of reaching a 76% MTY indigoidine from *p-*CA was achievable with 10 rounds of genetic modifications: 6 deletions predicted across 2 rounds of the PSP workflow, 3 targets from rational strain design, and ALE (**Figure 5C**).

With a highly curated model or strong mechanistic understanding, predictions can be translated into complex strain designs. Our previous study on growth coupling with glucose using multiplex CRISPRi led to ∼45% indigoidine MTY without the need for ALE (Banerjee *et al*, 2020). For these *p-*CA growth coupling sets we encountered many technical issues, including failure of gRNA-mediated knockdown with the CRISPRi approach used initially and secondly, synthetic lethality when a partial cut set was generated in the form of a triple deletion strain. While in this case the computational model cannot suggest a path forward beyond metabolism, our experience in strain engineering suggested that a partial cut set implementation followed by ALE was the best approach forward. ALE has been long recognized for its potential to generate gain of function mutations relevant for growth coupling (Fong *et al*, 2005), especially in the context of substrate catabolism (Van Hofwegen *et al*, 2016; Lim *et al*, 2021; Gao *et al*, 2022). Our predictions initially indicated considerable growth defects in these strains (**Figure 1A**) but we isolated strains after ALE with comparable growth and indigoidine titers to the WT control. Furthermore we did not observe growth defects in our Design 1 strains until the very last iteration with 10 total edits. Even partial growth coupled designs may impose an equally challenging (and distinct) evolutionary landscape, owing to the challenges posed when key metabolic pathways are removed. The gain of function mutants identified by whole genome sequencing likely reflect the evolutionary path (Bajić *et al*, 2018) our base strain navigated to restore growth on *p-*CA, in contrast to WT *P. putida* used previously (Mohamed *et al*, 2020). The role of cell shape and structure as well as the underlying molecular mechanisms for their applicability to strain engineering are only beginning to come into focus (Liu *et al*, 2022).

The utility of proteomics datasets to build a process understanding cannot be overstated. Proteomics analysis identified that oxidative stress was likely a key cause of inviability in the pre- ALE Δ3 strain, which was alleviated by the upregulation of the Dyp and AhpCF peroxidases in the Design 1 isolates. It also identified FleQ, a master regulator of chemotaxis that could be deleted, considerably increasing indigoidine titers, and was crucial to reweighting the metabolic model for the third and final DBTL cycle employed. Proteomics information sits in the “sweet spot” for characterizing cellular processes; it provides a rapid, semi-quantitative characterization of many activities much wider than metabolomics and can more directly implicate the cellular effectors than whole genome resequencing. Moreover, our analysis pipelines are generalizable as preparing cells for proteomic analysis is an accessible method for hard to lyse microbes like Corynebacteria or filamentous yeasts, where rapid metabolite quenching is challenging (Wellerdiek *et al*, 2009; Khanijou *et al*, 2022).

In conclusion, the multiple iterations represent four DBTL cycles in this study that achieved high TRY starting with predictions generated with a GSMM and the generation of a triple deletion mutant strain. This strain in turn was subject to ALE to address aspects outside of the model such as oxidative stress, chemotaxis, and inefficient substrate catabolism. The final strain produces 7.3 g/L indigoidine at 77% MTY in M9 *p-*CA minimal salt medium. Aided by rapid strain engineering we implemented learnings from a comprehensive cellular analysis to maximize TRY. This approach is likely generalizable to other microbes, carbon streams, and final products.

## Materials and Methods

*In silico* methods and *P. putida* genome reassembly

### Computation of constrained minimal cut sets

*Pseudomonas putida* KT2440 genome scale metabolic model (GSMM) iJN1411 and iJN1463 (Nogales *et al*, 2020) were used. Aerobic conditions with glucose as the sole carbon source were used to model growth parameters. The ATP maintenance demand and *para*-coumarate (*p*-CA) uptake were 0.97 mmol ATP/gDW/h and -10 mmol *p*-CA/gDW/h, respectively. Constrained minimal cut sets (cMCS) were calculated using the MCS algorithm (von Kamp & Klamt, 2017) available as part of CellNetAnalyzer (Klamt *et al*, 2007). Excretion of byproducts was initially set to zero, except for the reported secreted products specific to *P. putida* (gluconate, 2- ketogluconate, 3-oxoadipate, catechol, lactate, ethanol, methanol, CO2, and acetate). We calculated the maximum theoretical yields (MTY) for indigoidine using *p*-CA as the carbon source and the heterologous 2 gene indigoidine production pathway. Additional inputs including minimum demanded product yield, YPS (10% to 85% of MTY) and maximum demanded biomass yield, YBS, at 10 to 25% of maximum biomass yield were also specified in order to constrain the desired design space. The maximum size of cMCS was kept at the default (i.e., 50 metabolic reactions). Knockouts of export reactions and spontaneous reactions were not allowed. With the specifications used herein, each calculated knockout strategy (cMCS) demands production of indigoidine even when cells do not grow. Each cMCS computation run provides a prediction of several cMCS designs of the same cMCS size, but the cMCS sizes may vary depending on the additional inputs (YPS and YBS) for each run. All cMCS calculations were done using API functions of CellNetAnalyzer (Klamt *et al*, 2007) on MATLAB 2017b platform using CPLEX 12.8 as the MILP solver. The different runs, respective number of cut sets and number of targeted reactions to satisfy coupling constraints are included in **Supplementary Table 3**.

### Computation of elementary modes

For elementary modes analysis, a small model representing the central carbon metabolism of *P. putida* and the heterologous indigoidine production pathway (**Supplementary Data 3**) was used to calculate elementary modes by *efmtool* (Terzer & Stelling, 2008).

### Validation of constrained cut sets with constraint-based methods

*P. putida* GSMM was extended to account for indigoidine biosynthesis pathway and checked for strong growth coupling to confirm the chosen engineering strategy for experimental implementation. The cytosolic reaction for indigoidine biosynthesis from glutamine was as previously described (Banerjee *et al*, 2020). Flux balance analysis (FBA) was used to calculate the maximum theoretical yield (MTY) from reaction stoichiometry and redox balance. Flux variability analysis (FVA) was used along with FBA to check for minimum and maximum glutamine or indigoidine flux under the identified cMCS strategy to confirm growth coupling. FVA was performed with maximization of biomass formation as the objective function and the proposed gene deletions in each cMCS strategy along with constraints that were used for cMCS calculations. A positive minimum and maximum flux through the exchange reaction for the metabolite of interest (glutamine or indigoidine) confirmed growth coupling. COBRA Toolbox v.3.0 (Heirendt *et al*, 2019) in MATLAB R2017b was used for FBA and FVA simulations with the GLPK (https://gnu.org/software/glpk) or Gurobi Optimizer 8.1 (http://www.gurobi.com/) as the linear optimization solver.

### Proteomics data integration to create context specific models

The proteomics data and the *P. putida* GSMM, iJN1463, were used to extract context specific models for the two Design 1 isolates. We used the integrated metabolic analysis tool (iMAT) algorithm (Zur *et al*, 2010), which uses discrete levels for expression values: low, medium and high. iMAT allowed the integration of proteomics data into the GSMM and maximized highly and minimized lowly expressed reactions from the model. We used the log2 fold changes in protein counts with respect to WT in the two D1 isolates to perform this classification with a cutoff of half a standard deviation above the mean of the log2FC values, for active reactions, and half a standard deviation below for inactive reactions. The iMAT algorithm iteratively searches for a set of active reactions, within the cutoff range, linked to the levels of the associated proteins that simultaneously are able to result in biomass and indigoidine production. Simulations were performed using the iMAT implementation in the Cobra Toolbox with Matlab 2017b (Mathworks Inc., Natick, MA, USA) and CPLEX 12.8 as the solver.

### Nanopore + Illumina hybrid method DNA assembly, SNP analysis

*P. putida* genomic DNA of the indicated genotypes was prepared for whole genome sequencing and hybrid assembly using conventional laboratory techniques as previously described (Langley *et al*, 2019). Briefly, cryostocks were struck to single colonies on LB agar medium and inoculated into liquid 5 mL LB cultures and incubated overnight at 30 °C until saturated. Approximately 1.5 mL of cells were harvested, spun down (5,000 x *g*, 3 minutes), flash frozen and lysed (50 U/mL units RNAse, 0.1% w/v SDS, 200mM NaCl, pH 8) for genomic DNA extraction using 1.5 mL of cell culture with a phenol chloroform extraction and phase lock tubes (Qiagen Sciences, Germantown, MD) to enhance phase separation. Following isopropanol + NaOAc precipitation, the DNA pellet was washed twice with 70% ethanol and air dried before resuspension. Approximately 2 µg of DNA was used for either Illumina or Nanopore sequencing, conducted by SeqCenter Inc. (Pittsburgh, PA). For Illumina short read sequencing, sample libraries were prepared using the Illumina DNA Prep kit and IDT 10bp UDI indices and sequenced on an Illumina NextSeq 2000, producing 2x151bp reads. Demultiplexing, quality control and adapter trimming was performed with bcl-convert (v3.9.3) (Illumina Inc). Oxford Nanopore reads were processed for quality control and adapter trimming using porechop (github.com/rrwick/Porechop). Hybrid assembly with Illumina and ONT reads was performed with Unicycler (Wick *et al*, 2017). Assembly statistics were recorded with QUAST. Annotations of protein-encoding open reading frames and noncoding RNAs (ncRNAs) were predicted with the NCBI Prokaryotic Genome Annotation Pipeline (Tatusova *et al*, 2016). To map existing GeneID accession numbers (PP_XXXX) to the new chromosomal assemblies, we used the Geneious WGS software package (v10.0.1, BioMatters LTD). The new chromosomal assembly was aligned with the NCBI reference sequence (NC_002947.4) using Mauve and the transferred gene annotations were extracted as LCB (locally colinear blocks) alignment files. Single nucleotide polymorphisms and other variants were identified using the Geneious “Variations/SNP” (default settings for variant probability; bacterial translation start codon settings) tool comparing the mutant strains to either the WT strain or the pre-ALE Δ3 strain as indicated. Pre-existing mutations in the genome were deduced and excluded from further analysis by filtering existing mutations in our resequenced wildtype *P. putida* KT2440 as well as the pre-ALE Δ3 strain.

### Reagents and Bacterial Cultivation Conditions

#### Molecular Biology

All strains and plasmids used in this study are described in **Supplementary Table 6**. Cloning of synthetic DNA constructs was conducted using chemically competent DH10-β cells purchased from NEB (New England Biolabs, Ipswitch, MA) or XL-1 Blue (Agilent Technologies, Santa Clara, CA)cells prepared for chemical competency using the Inuoe method (Sambrook & Russell, 2001) by the UC Berkeley QB3 Core facility (Berkeley, CA). Two constructs used for the initial multiplex CRISPRi/dCpf1 prototyping were synthesized by Genscript USA Inc (Piscataway, NJ) using the same logic for unique promoter-gRNA-terminator designs as described previously (Banerjee *et al*, 2020). All DNA assemblies were designed using Snapgene (BioMatters Ltd). NEB OneTaq 2x PCR Master Mix was used for routine genotyping of *P. putida* transformants after recombineering or allelic exchange, and NEB Q5 2x PCR master mix was used to amplify DNA fragments for isothermal HiFi assembly (NEB). PCR and isothermal assembly were conducted following manufacturer’s guidelines for extension time and annealing temperature. The annealing time was always set to 3 seconds (Mamedov *et al*, 2008). Plasmids were transformed into *P. putida* strains of the indicated genotype via electroporation after 2 washes in 10% glycerol exactly as previously described (Banerjee *et al*, 2020). Both *E. coli* and *P. putida* glycerol stocks were stored at -80 °C using a final glycerol concentration of 25% (w/v). New gRNA designs were verified by Sanger sequencing (Azenta Life Sciences, Burlington, MA). Complex new DNA assemblies for heterologous gene pathways were verified by whole plasmid sequencing (Primordium Labs, Monrovia, CA). Routine growth of either microbe was carried out using Luria Bertani medium (LB, 10 g/L tryptone, 5 g/L yeast extract, 10 g/L NaCl) from Millipore Sigma. Kanamycin (50 µg/mL) or gentamicin (30 µg/mL) was added to the appropriate medium as indicated for experiments requiring the selection of plasmids. To assess strain viability using semi-quantitative serial dilution assays, strains were diluted 10x in a microtiter dish and 3 µL of diluted cells were plated on solid agar plates supplemented with LB and M9 media with additives as indicated and incubated at 30 °C. LB plates were photographed after 24 hours. M9 *p-*CA plates were photographed after 48 hours.

### Construction of targeted genomic mutants in *P. putida* via allelic exchange or recombineering

Generation of in-frame precise deletions in *P. putida* was implemented primarily using a recombineering/CRISPR counterselection protocol (Czajka *et al*, 2022). In brief, 90 bp ssDNA oligos were designed to bind 45 bp upstream and downstream precisely targeting the start and stop codons of the desired gene for removal. If the reading frame overlapped with other coding sequences, the deleted region was modified to exclude any bases that were necessary for expressing the overlapping coding sequences. All oligos were synthesized by IDT (Integrated DNA Technologies, Redwood City, CA) and are described in **Supplementary Table 7**. Recombineering clones were selected using Cpf1-based counterselection with a 20-21 bp gRNA with homology for the WT locus of the targeted gene for deletion. gRNAs were designed by identifying a 5’-TTTN-3’ PAM sequence 5’ upstream of the locus targeted for recombineering. All gRNA sequences are described in **Supplementary Table 8**. Cpf1/Cas12a endonuclease eliminates unedited cells by targeting the specified WT locus to generate a lethal double-strand break. After genotyping colonies by colony PCR and loss of the Cpf1-gRNA plasmid the process was repeated 2 more times. The inducer, 3-methyl-benzoate (3MB, m-toluic acid, Sigma T36609), was used at a concentration of 1mM. The 500 mM 3MB stock solution was dissolved in 50% ethanol/water, stored at 4 °C, and discarded after 6 months. Gene deletions were confirmed by colony PCR genotyping using NEB Onetaq Quick-Load 2X Master Mix with Standard Buffer (Cat. No. M0486L) as described above following an initial colony boiling step (50 µL 20mM NaOH, 30 minutes, 98 °C).

Several allelic exchange plasmids were also used for the integration of the original or variant heterologous indigoidine pathway, and for the deletion of PP_4120. Standard protocols for the initial integration, counterselection, and verification of strains by genotyping were executed without modification as previously described (Eng *et al*, 2021).

### Preparation of stock *p*-CA

A solution of *para*-coumarate (*p-*CA, 98% purity, Sigma Aldrich, Cat. No. C9008) was prepared at a concentration of 0.5 M by dissolving 20.52 g into 250 mL of sterile ddH20 and pH adjusted to 8 using NaOH pellets (Cat. No. S8045) with stirring to ensure complete dissolution. Afterwards, the solution was filter sterilized (0.2 µm SFC, Nalgene, Cat. No. 291-4520) and was discarded roughly after 3 months of storage at room temperature when the color changed from slightly amber to a dark brown. The rate of dark color appearance was not dependent on storage temperature, time of year, or exposure to ambient light, but may correlate with the age of specific batches purchased from the supplier.

### Cultivation of *P. putida* for growth and production of indigoidine

An overnight LB liquid culture was prepared from a single colony struck out from cryostorage. M9 minimal medium used here was modified from the “NREL” formulation (Linger *et al*, 2014) to 100 mM NH4Cl, 47.9 mM Na2HPO4, 22 mM KH2PO4, 8.56 mM NaCl, 2 mM MgSO4, 100 µM CaCl2 with 1X trace metals solution (Catalog Num. T1001, Teknova Inc, Hollister CA), 60 mM of *p-*CA and 30 mM of MOPS (pH 7.0, Sigma Catalog Num. M1254) was used to test the performance of the strains, or with *p*- CA concentrations as otherwise indicated for C/N ratio analysis. To inoculate cells for adaptation, 500 µL (starting OD600= 0.4) from a saturated LB overnight culture were added to a 5 mL of M9 *p-*CA medium in a culture tube. Cultures were incubated at 30 °C with 200 rpm with orbital shaking and 40% humidity. A second adaptation was performed in M9 medium in the desired format for the production run. Unless otherwise indicated, the concentration of *p-*CA used for indigoidine production was 60 mM. For when dCpf1/CRISPR plasmids were used for gene interference (CRISPRi) with Design I and Design II plasmids, 500 µM IPTG and 50 µg/mL kanamycin was added to the medium to induce the gRNA array and to maintain plasmid selection, respectively. IPTG was added at the same time the pathway was induced with arabinose. The majority of experiments here using the Design 1 strains did not require IPTG addition or plasmid selection during the indigoidine production runs with the exception of the proteomics data described in Figure 3B which used kanamycin for plasmid selection of the kanR+ pTE449 empty vector transformed into all 3 strains. When cultivated in a 24-deep well plate (Axygen Scientific, Union City, CA) a 1.5 mL volume fill was used and the experiment was done in quadruplicates with linear shaking at 1,000 rpm, 30 °C and 70% humidity using a gas-permeable film (114 µm AeraSeal Film, Omega Biotek, Cat. No. AC1201-02). For shake flask experiments, 20 mL or 50 mL cultures were grown in 250 mL Erlenmeyer baffled shake flasks with orbital shaking at 200 rpm in triplicates. Indigoidine pathway gene expression was induced with 0.3% w/v or 1.5% w/v arabinose as indicated in the figure legend. Error bars indicate the standard deviation between measurements.

### Kinetic Growth Curves

A kinetic time course to monitor changes in optical density was used to measure at OD595 nm every 15 minutes using a Molecular Devices Filtermax F5 plate reader (Molecular Devices LLC, San Jose, CA) set to “high” shake speed and linear shaking mode in a 24 well microtiter dish plate sealed with a Breathe-Easy^Ⓡ^ transparent membrane (Sigma-Aldrich). For the data in **Supplementary Figure 9**, a Molecular Devices Spectramax 5e was used to collect the timecourse by monitoring OD600 with the same plate format but sampled every 10 minutes for the duration of the timecourse. Microtiter plates were incubated at 30 °C for the duration of the timecourse.

### Glassware

Our facility shares core media preparation and glasswash services with fungal and plant biology research groups without standardized glassware cleaning protocols across organisms. *P. putida* is extremely sensitive to residual detergents (ie Alconox, Dove, “Boardwalk Pots and Pans”) which can also compromise strain growth and final product titers. Glassware was inspected before use for residual powder, debris or unusual bubbles forming after being filled with culture medium. As needed, glassware was recleaned by triple rinsing with MilliQ grade ddH2O and 95% ethanol, which was repeated six times. Glassware was then resterilized by autoclaving.

### Adaptive laboratory evolution

We implemented adaptive laboratory evolution (ALE) using several different concurrent strategies and independent strain isolates. The goal of this ALE experiment was to restore growth of the engineered strains in M9 *p-*CA with *p-*CA concentrations exceeding 100 mM without a pre-determined maximum *p-*CA concentration at the start of the experiment. These regimes included titrating the ratio of LB to 1x M9 *p-*CA medium initially favoring 100% LB and serially passaging saturated cultures to fresh 80% LB medium containing 20% 1x M9 10 mM *p-*CA medium. Cells were grown aerobically with shaking at 30 °C. We gradually increased the percentage of M9 *p-*CA in dilute LB medium. We also tried first growing cells in M9 medium containing both glucose and *p-*CA where the *p-*CA concentration was gradually increased. The third method attempted was by plating cells directly on M9 60 mM *p*-CA plates from saturated overnight LB cultures. The Δ3 strain, the Δ3 Δ*gacA* strain, and a Δ4 (ΔPP_0897 ΔPP_1755 ΔPP_0944 ΔPP_1378) strain were all tested with this passaging regimen with at least 3 biological replicates of each genotype. These strains used and how they were constructed are diagrammed in **Supplementary Figure 2**. As arabinose was present in the culture medium, we visually inspected cultures over the duration of this experiment to determine if they still turned blue as a crude indicator of an active indigoidine pathway. Blue forming cultures were only recovered in the Δ3 strain with *p-*CA as the sole carbon source, but not from cultures that also contained glucose. Some clones were recovered in the Δ3 Δ*gacA* background but no longer were blue colored. Many independent lineages were lost upon serial passage and could not be recovered. Other lineages failed to revive on LB plates after short term storage as cryostocks at -80 °C.

### Peroxide Sensitivity Assay

*P. putida* sensitivity to exogenously added hydrogen peroxide was assayed as previously described (Nikel *et al*, 2021; Jensen *et al*, 2017) with slight modifications. Briefly, single colonies of the appropriate genotype were struck out from glycerol stocks and inoculated into 5 mL LB cultures and incubated with shaking at 30 °C overnight to generate saturated cultures. 3 µL of these cultures were used to inoculate replicate wells in a 48 round well microtiter plate, where each well was prefilled with 300 µL LB medium. Hydrogen peroxide was freshly added to aliquots of LB medium immediately before the plate was prepared at concentrations of 0 mM, 3 mM, and 10 mM from a 30% w/v stock hydrogen peroxide solution (Sigma Aldrich). The results plotted are the average value of three biological replicates, and the banded shaded area represents the standard deviation of the measurements. All experiments were repeated at least one additional time on a different day.

### Microbial Interaction Assay

*P. putida* WT, Design 1b, and the pre-ALE Δ3 strain were tested for their ability to restore growth in a *P. putida* Δ*fcs* strain using an established microbial interaction assay (Eng *et al*, 2020). Briefly, biomass was first spread from a Δ*fcs* strain in a thick horizontal line down the center of an M9 60 mM *p-*CA agar plate; the potential complementing strain was next spread in a vertical line, intersecting the Δ*fcs* strain in the middle. The plates were incubated at 30 °C for 10 days and visually inspected for biomass formation in the Δ*fcs* strain each day.

### Soft X-Ray Tomography Characterization of Cells

*P. putida* WT and Design 1a cells were prepared for soft X-ray tomography (SXT) as follows. Briefly, cells were grown either in rich LB medium or adapted for growth in M9 60 mM *p-*CA medium. Several hours before cells were to be prepared for SXT, an additional 20% initial volume of fresh medium was added to the cultures to restore consumed nutrients in preparation for transport. Cultures were transferred out of culture tubes and into 50 mL falcon tubes for transit to the National Center for X-ray Tomography (NCXT) for cryo-fixation before SXT imaging. Upon arrival, cells were prepared for cryofixation via loading 1-2 microliters into pre-cut tapered glass capillaries (∼6um opening) using micro-loaders (Chen *et al*, 2022a). Briefly, the concentration of *P. putida* cells was analyzed using a TC20 Automated Cell Counter (Bio-Rad Laboratories, Hercules, CA). Cells were then harvested by centrifugation at 8000 rpm for 3 minutes and loaded into capillary tubes using a micro-tip. The prepared specimens in capillary tubes were then plunged frozen into liquid nitrogen cooled liquid propane and then stored in a LN2 Dewar until SXT imaging. Prior to data acquisition, samples were loaded onto a transmission soft X-ray microscope using home-built cryo-transfer apparatus to ensure the integrity of vitrified samples. Data acquisition for 3D tomographic reconstruction was done by taking 2D projections from different angles. The equipped full rotation sample stage rotates with 2-degree increments for 92 projections to ensure an isotropic image reconstruction (Parkinson *et al*, 2012). SXT data reconstruction without fiducial markers was done by implementing AREC3D (Parkinson *et al*, 2012), and the auto-segmentation pipeline was based on the CNN training (Ekman *et al*, 2020). Cell size and density measurements were manually quantified using the Fiji software package and Amira. At least 100 cells were quantified per timepoint and genotype. Statistical analysis was performed using a two-tailed unpaired *t*-test for the volumetric and density comparisons indicated in the figure to determine if the changes were statistically significant.

### Shotgun Proteomics Analysis

The Design 1a and Design 1b strains and a WT control strain transformed with an empty vector pTE449 plasmid were grown in triplicates in M9 60 mM *p*-CA 50 µg/mL kanamycin using 250 mL shake flasks and back diluted to a starting OD of 0.05 OD and 8 hours post back dilution the samples were harvested when cells were in mid-log phase and stored at -80 °C until sample preparation. For the double deletion strain analysis, all combination of double deletion strains and the D1b strain (ΔPP_1378, ΔPP_1755, ΔPP_0944) were grown in quadruplicates in M9 60 mM *p-*CA media using the 24 well deep well plate format and back diluted to a starting OD of 0.05 and allowed to grow for 8 hours until the cell OD reached 0.06 and similarly harvested and stored at -80 °C until analysis. After all samples were collected, protein was extracted from harvested *P. putida* strain cultures and tryptic peptides were prepared by following established proteomic sample preparation procedures (Chen *et al*, 2023). Briefly, cell pellets were resuspended in Qiagen P2 Lysis Buffer (Qiagen Sciences, Germantown, MD, Cat.#19052) to promote cell lysis. Proteins were precipitated with addition of 1 mM NaCl and 4 x volume acetone, followed by two additional washes with 80% acetone in water. The recovered protein pellet was homogenized by pipetting mixing with 100 mM Ammonium bicarbonate in 20% Methanol. Protein concentration was determined by the DC protein assay (BioRad Inc, Hercules, CA). Protein reduction was accomplished using 5 mM tris 2-(carboxyethyl)phosphine (TCEP) for 30 minutes at room temperature, and alkylation was performed with 10 mM iodoacetamide (IAM; final concentration) for 30 minutes at room temperature in the dark. Overnight digestion with trypsin was accomplished with a 1:50 trypsin:total protein ratio. The resulting peptide samples were analyzed on an Agilent 1290 UHPLC system coupled to a Thermo Scientific Orbitrap Exploris 480 mass spectrometer for discovery proteomics (Chen *et al*, 2022b). Briefly, peptide samples were loaded onto an Ascentis® ES-C18 Column (Sigma–Aldrich, St. Louis, MO) and separated with a 10 minute LC gradient (10% Buffer A (0.1% FA in water) – 35% Buffer B (0.1% FA in ACN)). Eluting peptides were introduced to the mass spectrometer operating in positive-ion mode and were measured in data-independent acquisition (DIA) mode with a duty cycle of 3 survey scans from m/z 380 to m/z 985 and 45 MS2 scans with precursor isolation width of 13.5 m/z to cover the mass range. DIA raw data files were analyzed by an integrated software suite DIA-NN (Demichev *et al*, 2020). The database used in the DIA-NN search (library-free mode) is the latest Uniprot *P. putida* KT2440 proteome FASTA sequence plus the protein sequences of heterogeneous pathway genes and common proteomic contaminants. DIA-NN determines mass tolerances automatically based on first pass analysis of the samples with automated determination of optimal mass accuracies. The retention time extraction window was determined individually for all MS runs analyzed via the automated optimization procedure implemented in DIA-NN. Protein inference was enabled, and the quantification strategy was set to Robust LC = High Accuracy. Output main DIA-NN reports were filtered with a global FDR = 0.01 on both the precursor level and protein group level. A jupyter notebook written in Python executed label-free quantification (LFQ) data analysis on the DIA-NN peptide quantification report, and the details of the analysis were described in the established protocol (Chen & Petzold, 2022).

### TCA Metabolite Targeted Metabolomics analysis of growth coupled strains

Organic acids were measured via reversed-phase chromatography and high-resolution mass spectrometry. Liquid chromatography (LC) was conducted on an Ascentis Express RP-Amide column (150-mm length, 4.6-mm internal diameter, and 2.7-μm particle size; Supelco, Sigma-Aldrich, St. Louis, MO, USA), equipped with the appropriate guard column, using an Agilent Technologies 1260 Series high performance liquid chromatography (HPLC) system (Agilent Technologies, Santa Clara, CA, USA). A sample injection volume of 2 μL was used throughout. The sample tray and column compartment were set to 5 °C and 50 °C, respectively. The mobile phase solvents used in this study were of LC-MS grade and purchased from Honeywell Burdick & Jackson (Honeywell, Charlotte, North Carolina, USA). Other chemicals and reagents were purchased from Sigma-Aldrich. The mobile phases were composed of 0.1% formic acid/84.9% water/15% methanol (v/v/v) (A) and 0.04% formic acid and 5 mM ammonium acetate in methanol (B). Organic acids were separated via gradient elution under the following conditions: linearly increased from 0% B to 30% B in 5.5 minutes, increased to 100% B in 0.2 minutes, held at 100% B for 2 minutes, decreased from 100%B to 0%B in 0.2 minutes, and held at 0%B for 2.5 minutes. The flow rate was held at 0.4 mL/minute for 7.7 minutes, linearly increased from 0.4 mL/minute to 1 mL/minute in 0.2 minute, and held at 1 mL/minute for 2.5 minutes. The total LC run time was 10.4 minutes. The HPLC system was coupled to an Agilent Technologies 6545 series quadrupole time-of-flight mass spectrometer (QTOF-MS). MS conditions were as follows: Drying and nebulizing gases were set to 10 L/minute and 25 lb/in2, respectively, and a drying-gas temperature of 300 °C was used throughout. Sheath gas temperature and flow rate were 330 °C and 12 L/minute, respectively. Electrospray ionization, via the Agilent Technologies Jet Stream Source, was conducted in the negative ion mode and a capillary voltage of 3,500 V was utilized. The fragmentor, skimmer, and OCT 1 RF Vpp voltages were set to 100, 50, and 300 V, respectively. The acquisition range was from 70-1,100 m/z, and the acquisition rate was 1 spectra/sec. The QTOF-MS system was tuned with the Agilent ESI-L Low concentration tuning mix (diluted 10-fold in a solvent mixture of 80% acetonitrile and 20% water) in the range of 50-1700 m/z. Reference mass correction was performed with the Agilent Technologies API-TOF Reference Mass Solution Kit. Data processing and analysis were conducted via Agilent Technologies MassHunter Qualitative Analysis, Profinder, and/or Quantitative Analysis. All other metabolites were analyzed according to the method described by Amer et al. (Amer *et al*, 2022). Metabolites were quantified via seven-point calibration curves from 0.39 to 25 µM. To calculate statistical significance, we used an unpaired two tailed *t*-test to compare concentrations between the WT control strain and the D1b *glnA* Δ*fleQ* strain. Exact *p* values are reported.

### Analytical Methods for the Quantification of Aromatics

All samples were stored at -20 °C except unless otherwise mentioned. *p*-CA consumption was analyzed by High Performance Liquid Chromatography as previously described (Rodriguez *et al*, 2019). Briefly, we used an Eclipse Plus Phenyl-Hexyl column (250 mm length, 4.6 mm diameter, 5 µm particle size, Agilent Technologies) and a Diode Array Detector (G4212, Agilent Technologies) to measure UV absorption at 254 nm, 310 nm, and 280 nm. Cultures were centrifuged at 6010 x *g* to remove cells and the extracellular supernatants were harvested. After all cultures were processed, the supernatant was diluted with mobile phase (10mM Ammonium acetate 0.03% formic acid) and filtered using a 0.45 µm 96-well microplate (polypropylene membrane, Cat. No. 200983-100, Agilent) by centrifugation for 30 minutes at 5,250 x *g*. Filtrates were collected in a 96-well plate sealed with an EZ-Pierce film (Cat. No. Z721581, Sigma Aldrich) for analysis by HPLC. Fresh *p-* CA standards were prepared on each day to quantitate *p-*CA concentration in the samples. Finally, phenolic compounds were also analyzed according to the LC-MS method as previously described (Eudes *et al*, 2013).

### Purification of indigoidine used for product quantification and verification of purity by NMR

*Pseudomonas putida* KT2440 harboring an integrated indigoidine expression cassette (“WT Control”, or “Control strain”) was used to produce indigoidine which was then used for purification and generation of a standard curve for colorimetric analysis as previously described (Banerjee *et al*, 2020) with slight modifications. Batch to batch variation in commercially procured *p-*CA (Sigma-Aldrich) impacted downstream processing and extraction. Triplicate 50 mL liquid cultures were prepared in baffled shake flasks as described above using M9 60 mM *p-*CA and 1.5% (w/v) arabinose. The cell pellets were harvested after 24 h and washed with a series of solvents from higher to lower polarity for the purification of indigoidine (water, methanol, ethyl acetate, acetonitrile, acetone and hexane). Each wash step consisted of solvent addition, vortexing for ∼1 minute and centrifuging at 5,250 x *g* for 5 minutes to recover indigoidine in the insoluble fraction. Each solvent wash was repeated twice. The resulting product was dried overnight to evaporate residual hexane and the following morning was ground to a fine powder. Purity was confirmed by 1H-Nuclear magnetic resonance (NMR) as described previously (Banerjee *et al*, 2020).

A standard curve was generated using purified indigoidine generated above. The scale used was calibrated to 25 mg using calibration weights (Troemner, ASTM Class 4) and to account for measurement error. 25 mg aliquots of purified indigoidine were mixed with 25 mL of 100% DMSO in a 50 mL falcon tube and incubated overnight on a shaking platform covered with aluminum foil to ensure total indigoidine solubilization. The reference samples were prepared in triplicate following the triplicate indigoidine samples purified above. Indigoidine absorbance was measured at 612 nm (Wehrs *et al*, 2019a) using 1.25-fold serial dilutions of the prepared stock solution to correlate OD612 to mg/L indigoidine, and a formula was generated. The formula was : Y= 0.2933x- 0.013 and was in good agreement with our previously reported formulae for both *P. putida* and *E. coli* prepared samples from glucose bioconversion processes.

### Colorimetric determination of indigoidine from crude sample extracts

200 µL samples were harvested at the designated timepoints and stored at 4 °C until samples were batch processed, for no longer than 24 hours. When ready, samples were pelleted at 6,010 x g for 3 minutes using a bench top centrifuge (Eppendorf 5424/5424R). 100% DMSO was added to resuspend the cell pellet with gentle pipetting until the pellet was completely broken down. DMSO resuspended cell pellets were vortexed for 45 minutes in a Plate Incubator (Cat. No. H6004, Benchmark scientific). Once extraction was completed, the samples were centrifuged at 21,130 x *g* for 1.5 minutes to remove contaminating cell debris. 100 µL of each sample was added to a 96-well clear flat-bottom plate (Cat. No. 353072, Corning Incorporated) to read absorbance at 612 nm using a microplate reader (SpectraMax M2, Molecular Devices). The standard curve was applied to convert OD612 readings into indigoidine titers (g/L). The M9 medium control and WT *P. putida* cell mass do not contribute appreciable signal to the DMSO OD612 readings.

### LC-MS-Based Methods for the Quantification of Indigoidine

1.5 mL were sampled from liquid cultures in M9 *p*-CA at the 24 h timepoint (or otherwise indicated) of an indigoidine production run and the extracted pellets were quenched with methanol by mixing thoroughly. The resulting lysate was mixed with water with a final concentration of 50% methanol and filtered through a 3,000Da MW/CO exclusion column (Amicon Ultra, Millipore, Cat. No. UFC500396) by centrifugation at 13,000 x *g* and -2 °C for 30-60 minutes to remove contaminating high molecular weight species.

Liquid chromatography (LC) was conducted on a Kinetex XB-C18 column (100-mm length, 3.0- mm internal diameter, and 2.6-μm particle size; Phenomenex, Torrance, CA USA) using a 1260 Series HPLC system (Agilent Technologies, Santa Clara, CA, USA). A sample injection volume of 3 μL was used. The sample tray and column compartment were set to 6 °C and 50 °C, respectively. The mobile phases were composed of 0.4% formic acid (Sigma-Aldrich, St. Louis, MO, USA) in water (A) and 0.4% formic acid in methanol (B). The mobile phase solvents used in this study were of LC-MS grade and purchased from Honeywell Burdick & Jackson (Honeywell, Charlotte, North Carolina, USA). Indigoidine was separated via gradient elution under the following conditions: linearly increased from 10% B to 74.4% B in 4.6 minutes, linearly increased from 74.4% to 90% in 0.4 minutes, held at 90%B for 1 minutes, linearly decreased from 90%B to 10%B in 0.2 minutes and held at 10%B for 2 minutes. The flow rate was held at 0.42 mL/minute for 5 minutes, linearly increased from 0.42 mL/minute to 0.65 mL/minute in 1 minute and held at 0.65 mL/minute for 2.2 minutes. The total LC run time was 8.2 minutes. The HPLC system was coupled to an Agilent Technologies 6520 series quadrupole time-of-flight mass spectrometer (QTOF-MS). Drying and nebulizing gases were set to 11 L/minute and 30 lb/in2, respectively, and a drying-gas temperature of 330°C was used throughout. Electrospray ionization was conducted in the negative ion mode and a capillary voltage of 3,500 V was utilized. The fragmentor, skimmer, and OCT 1 RF Vpp voltages were set to 140, 50, and 250 V, respectively. The acquisition range was from 70-1100 m/z, and the acquisition rate was 0.86 spectra/s. The QTOF-MS was tuned with the Agilent Technologies ESI-L Low concentration tuning mix in the range of 50-1700 m/z. Reference mass correction was performed with the Agilent Technologies API-TOF Reference Mass Solution Kit (part number G1969-85001) via a second ESI sprayer. Data acquisition and processing were conducted via the Agilent Technologies MassHunter Qualitative Analysis, Profinder, and/or Quantitative Analysis. Metabolites were quantified via a nine-point calibration curve from 1.95 to 500 µM.

### Generation of BCD Liquor of lignin from [Ch]Lys]-pretreated sorghum

Cholinium lysinate ([Ch][Lys])-based sorghum pretreatment and subsequent saccharification was performed in a one-pot configuration in a 1 L Parr 4520 series Bench Top reactor (Parr Instrument Company, Moline, IL) to obtain solid lignin residues as described previously (Choudhary *et al*, 2023). Briefly, 15 wt% 2 mm *Sorghum bicolor*, 10 wt% [Ch][Lys], and 75 wt% water were weighed in and pre- mixed in the Parr vessel to obtain a slurry. The obtained slurry was pretreated for 3 hours at 140 °C with stirring at 80 rpm powered by process (Parr Instrument Company, model: 4871, Moline, IL) and power controllers (Parr Instrument Company, model: 4875, Moline, IL) using three-arm, self-centering anchor with PTFE wiper blades. After 3 hours, the pretreated slurry was cooled down and pH was adjusted to 5 with concentrated hydrochloric acid. Enzymatic saccharification of the pH adjusted slurry was carried out at 50 °C for 72 hours at 80 rpm using enzyme mixtures Cellic® CTec3 and HTec3 (9:1 v/v) at a loading of 10 mg protein per g sorghum. After 72 hours, the slurry was centrifuged and washed multiple times with DI water to remove any residual sugars and [Ch][Lys]. The washed material was freeze dried to obtain [Ch][Lys]-Sorghum lignin and is fully characterized elsewhere (Wendt *et al*, manuscript in preparation). This lignin residue was used to generate BCD liquor in a 1 L Parr reactor. Lignin residue (wt%) and 5% sodium hydroxide (aqueous solution) were added to the Parr vessel and the temperature was ramped up to 120 °C in 30 minutes. The temperature was held at 120 °C for 30 minutes with vigorous stirring at 80% output of the rotor. After 30 minutes, the reactor was cooled down and pH was adjusted to 7 with 37% Hydrochloric acid and stored at 4 °C until used to replace all ddH2O in 1x M9 BCD liquor medium and sterilized by filter sterilization through a 0.2 µM filter. Major and minor aromatic constituents were characterized using LC-MS and are described in **Supplementary Table 5**. Since BCD liquor cannot be autoclaved for sterilization, we added 50 µg/mL chloramphenicol to the medium to inhibit growth of other microorganisms during indigoidine production runs. The resulting BCD liquor was used in place of ddH20 to make M9 minimal salt medium with BCD also substituting for all carbon present. The BCD liquor generation and indigoidine production run was repeated with a second batch of *Sorghum bicolor* and similar results were observed.

## ACKNOWLEDGEMENTS

We thank Jeff Czajka (PNNL), Hyungyu Lim (Inha University, South Korea), Adam Feist (UC San Diego), Corinne Scown (LBNL) and all members of the Mukhopadhyay group for insightful comments and suggestions throughout the formation of this manuscript. We also acknowledge Andrew Lau and Chunsheng Yan for technical assistance. TE, DB, JM, YC, HM, EB, RK, YLD, JG, CJP, BS, JDK, and AM are funded on the DOE - BER JBEI project, through contract DE- AC02-05CH11231 to the Regents of the University of California, who administers the subcontract between Lawrence Berkeley National Laboratory and the US Department of Energy. HC and JG are funded through Sandia National Laboratories, a multi-mission laboratory managed and operated by National Technology and Engineering Solutions of Sandia, LLC, a wholly owned subsidiary of Honeywell International Inc., for the U.S. Department of Energy’s National Nuclear Security Administration under contract DE-NA0003525. We thank Idaho National Labs (Idaho Falls, ID) for supplying the *Sorghum bicolor* used in this study. JHC, AE and CL are funded by the National Center for X-Ray Tomography and the Advanced Light Source, an Office of Science User Facility operated for the U.S. Department of Energy (DOE) Office of Science by Lawrence Berkeley National Laboratory, which was supported by the U.S. DOE under Contract No. DE- AC02-05CH11231. SXT data collection and analysis was partially supported by grant P30 GM138441 from the National Institute of General Medical Sciences of the National Institutes of Health.

## Author Contributions

Molecular biology and strain engineering: TE, JM, DB, AC. Computation of growth coupled designs and metabolic flux analysis: DB. Analytical Chemistry: EB, RK, YLD. Proteomics: YC, JG, CJP. X Ray Tomography: JHC, AE, CL. Generation of BCD liquor: HC, JG. Supervision, Formal Analysis, Acquisition of Funds: AM, JDK, BS and CJP. Drafted the manuscript: TE, DB, AM. All authors have read, provided feedback, and approved the manuscript for publication.

## DATA AVAILABILITY

Sequencing data for strains described in this study have been deposited at NCBI under BioProject ID No. PRJNA937293. The raw reads for strains sequenced with Illumina short read and Oxford Nanopore technology are deposited under BioSample No. SAMN33347499 - SAMN33347506. The assembled genomes have been deposited in the NCBI genome repository with BioSample No. SAMN33347507 - SAMN33347510 and their respective genome accession numbers CP118872 - CP118875. Proteomics data is available at PanoramaOnline at the following website link. Plasmid sequences and *P. putida* strains used in this manuscript have been deposited at the Joint BioEnergy Institute Strain Archive and can be requested at public-registry.jbei.org following execution of a Material Transfer Agreement. The generated mass spectrometry proteomics data have been deposited to the ProteomeXchange Consortium via the PRIDE (Perez-Riverol *et al*, 2022) partner repository with the dataset identifier PXD040697.

## CODE AVAILABILITY

All custom code used in this study is available in **Supplementary Data 4**.

